# Flavonoid chronotherapy normalizes circadian dynamics and sleep architecture in mice with painful nerve injury

**DOI:** 10.64898/2026.10.26.761906

**Authors:** H Launonen, W Dai, M Wen, N Milicevic, MK Koskinen, A Pertovaara, E Kalso, HK Wigren, V Palada

## Abstract

Neuropathic pain (NP) is linked to disrupted sleep and circadian abnormalities upon painful peripheral nerve injury. By performing spared nerve injury (SNI) in circadian reporter PER2::LUC mice, we characterized the real-time changes in rhythmicity of the core clock gene *Per2* in the pain-regulating dorsal root ganglia (DRG), and the sleep-wake cycle using wireless EEG/EMG under NP conditions. We further evaluated whether the clock-resynchronizing flavonoids, nobiletin (2 mg/kg/day) and tangeretin (1.8 mg/kg/day), administered continuously via osmotic minipumps for two weeks, could restore circadian dynamics and alleviate pain-related behaviour.

Both flavonoids alleviated SNI-induced dynamic hypersensitivity in PER2::LUC mice on post-surgical day 14. SNI increased the amplitude of *Per2* rhythms in the DRGs, which was restored to pre-surgical levels by administration of flavonoids. In addition, tangeretin showed sleep-restoring properties upon SNI by alleviating disturbances in NREM sleep and wakefulness. These results support targeting circadian disruption as a potential chronotherapeutic approach to alleviate NP.

## Introduction

Affecting over 500 million individuals globally^1^, neuropathic pain (NP) remains a major clinical challenge due to limited therapeutic efficacy and frequent co-occurrence with sleep disorders^2,3^. While traditional analgesic strategies focus primarily on pain transmission and modulation^4^, the broader systemic and physiological consequences, particularly the disruption of circadian dynamics, remain largely unaddressed by conventional pain management ^5–7^.

Circadian rhythms govern physiological processes across mammalian tissues. The master clock in the suprachiasmatic nucleus and peripheral tissue clocks coordinate circadian rhythms through clock genes, which regulate the rhythmic expression of genes involved in inflammatory, hormonal, and neurotransmitter signalling^5,8^. Emerging evidence indicates that pain is under circadian control^5,9^. The intensity of pain varies rhythmically across the day in various painful conditions^10^, being most severe in the evening in NP^7,11^. In addition, circadian disruption caused by night-shift work predisposes individuals to increased pain sensitivity^1^^2^. Addressing this, our recent work shows that repetitive circadian disruption via chronic jet lag worsens mechanical hypersensitivity in mice and profoundly alters daily transcriptional rhythms in spinal cord circuits following nerve injury^13^. Preclinical studies increasingly show that peripheral nerve injury disrupts the daily regulation of hundreds of genes^14^ in pain-regulating sensory tissues and reduces the spinal expression of the core clock gene *period 1* (*Per1)*^15^. While initial findings suggest the presence of circadian disruption in NP, this relationship remains largely unexplored.

Circadian disruption also extends to sleep. Most patients with NP report poor sleep quality, which can in turn reinforce pain^2^. Recently, we showed that rapid eye movement (REM) sleep is reduced in mice upon SNI^16^, a widely used and robust model of NP caused by ligation of two branches of the sciatic nerve while sparing the sural nerve^17^. Consistently, other animal models of NP also demonstrate marked alterations in sleep architecture^18–20^. At the molecular level, clock genes contribute to sleep regulation^21^. Period 2 (*Per2*), which controls the molecular clock by inhibiting CLOCK–BMAL1 activity^8^, is particularly relevant to sleep, as *Per2* gene expression is altered in response to sleep deprivation in the mouse brain^21,22^.

The observations linking NP to circadian abnormalities raise the possibility that the realignment of circadian rhythms could have therapeutic potential in NP. Nobiletin and its structural analogue tangeretin are natural citrus-derived polymethoxyflavones which have previously shown clock-modulating properties in fibroblasts and brain vessels of PER2::LUC mice^23–25^. Furthermore, nobiletin exerts analgesic effects in chronic constriction injury (CCI)-induced NP^26^ and endometriosis mouse model^27^. Both flavonoids have shown beneficial effects on sleep-related processes in mice, with nobiletin improving sleep consolidation in aged mice and tangeretin prolonging the duration of thiopental-induced sleep^28,29^. These properties make these two flavonoids interesting chronobiotic candidates for the treatment of NP.

Previous studies of circadian mechanisms in painful conditions have heavily relied on static expression measurements from tissues collected at several time points across the day, which cannot fully capture the dynamics of the molecular clock in real time^5,15^. By integrating SNI procedure with PER2::LUC mice, a commonly used luciferase knock-in reporter model of *Per2* rhythmicity^30^, we aimed to establish a new model that allows for real-time assessment of circadian abnormalities under NP conditions. Furthermore, we evaluated the flavonoids’ ability to restore circadian dynamics and alleviate pain, using osmotic pumps for continuous drug delivery, to minimise sleep-wake and circadian disturbances from repeated drug administration^16^. The changes in *Per2* rhythmicity in DRGs, containing nociceptors under circadian control^31^, were explored *ex vivo*. Moreover, the sleep-wake cycle in freely moving PER2::LUC mice with SNI or control sham surgery was monitored using wireless transmitters and we assessed the potential of flavonoids to restore sleep homeostasis upon painful nerve injury.

## Results

### Spared nerve injury causes mechanical and dynamic hypersensitivity in both sexes of PER2::LUC mice

To investigate circadian abnormalities in sleep architecture and in pain-processing tissue DRGs under NP conditions, PER2::LUC mice underwent wireless electroencephalography/electromyography (EEG/EMG) transmitter implantation (HDX02, Data Sciences International, USA) (Fig. 1A). Following a 14-day recovery period, vigilance states were recorded at baseline, and NP was induced in male and female mice by SNI surgery, in which two branches of the sciatic nerve were dissected, leaving the sural nerve intact^17^. Von Frey filament tests for static and brush tests for dynamic hypersensitivity were conducted at baseline before the surgeries and on post-surgical days 3, 7, and 14. EEG recordings were performed for 24 h at the baseline and post-surgical day 14, after which DRGs were dissected and placed into a LumiCycle 32 bioluminescence device (Actimetrics, Lafayette, IN, USA) for 5 days for real-time *Per2* rhythmicity monitoring.

**Figure 1.**
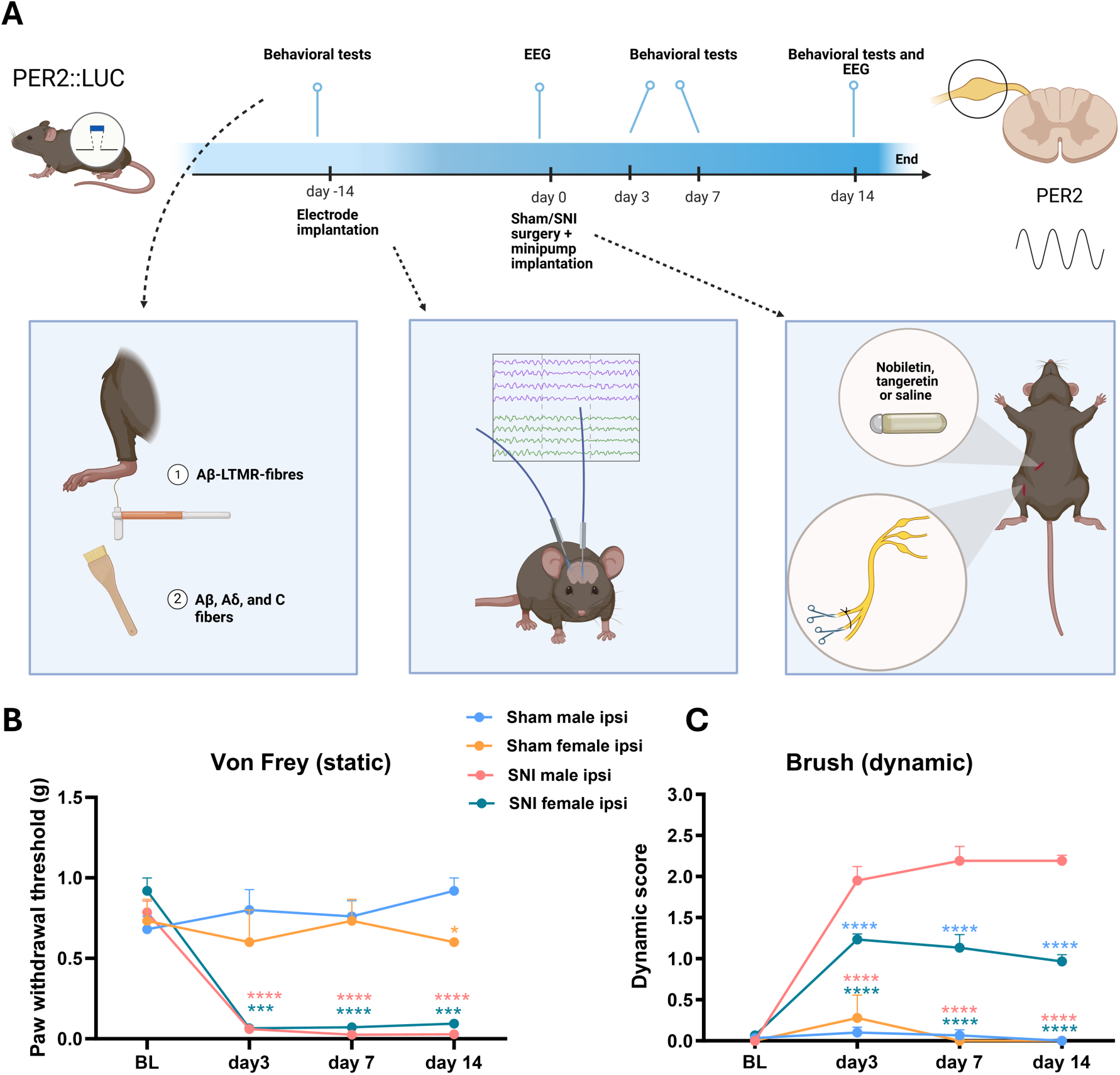
Study design and assessment of mechanical and dynamic hypersensitivity in sham- and spared nerve injury (SNI)-operated male and female PER2::LUC mice. **A)** To assess circadian abnormalities in the sleep-wake cycle and dorsal root ganglia under neuropathic pain (NP) conditions, PER2::LUC mice were equipped with electroencephalography/electromyography (EEG/EMG) transmitters. Following a two-week recovery period, baseline EEG measurements were recorder in freely moving mice before NP was established in both sexes using spared nerve injury (SNI) surgery. Mechanical pain sensitivity was assessed at baseline and on days 3, 7, and 14 after surgery using the von Frey filaments test, which sensitizes Aβ, Aδ, and C fibres, and the dynamic brush test, which sensitizes Aβ low-threshold mechanoreceptor (LTMR) fibers^50^. EEG/EMG was recorded again on post-surgical day 14, followed by DRG collection and five-day monitoring of PER2 rhythms using a Lumicycle 32 device. The PER2-luciferase fusion protein enables monitoring of PER2 oscillations by producing bioluminescence signal that reflects PER2 abundance over time. **B)** Paw withdrawal thresholds were significantly reduced in the operated hind paw of SNI mice compared with sham and remained decreased throughout the protocol. **C)** Brush test revealed increased responses following SNI, while responses in sham-operated mice remained at baseline in both sexes. The dynamic brush test was scored from 0 to 3, with 0 indicating no hypersensitivity and higher scores indicating gradually increasing hypersensitivity. Ipsi refers to the operated (ipsilateral) leg. EEG= electroencephalography, PER2= Period 2. Statistical analysis was performed using two-way ANOVA followed by Tukeýs post hoc test. Colored asterisks indicate statistically significant differences between the following groups: yellow, sham female ipsi *vs*. sham male ipsi; green, sham female ipsi *vs*. SNI female ipsi; red, sham male ipsi *vs*. SNI male ipsi; and blue, SNI male ipsi *vs*. SNI female ipsi. Values are expresses as mean ± SEM. ***p<0.001, ****p<0.0001. n=7 in the SNI male ipsi group, n= 5 in the sham male ipsi and SNI female ipsi groups, and n=3 in the sham female ipsi group. Figure 1A is generated using BioRender.

We first assessed the development of NP following SNI in male and female PER2::LUC mice. For von Frey responses, two-way ANOVA showed a significant time × group interaction (F(9, 64) = 12.2, p < 0.0001) (Fig. 1B). Painful nerve injury significantly decreased paw withdrawal responses in male and female SNI mice compared to sham mice starting from day 3 onward. In addition, female sham mice exhibited lower pain thresholds than sham males at day 14 (p = 0.037). Consistently, a significant time × group interaction was observed for dynamic brush responses (F(9, 64) = 21.5, p < 0.0001) (Fig. 1C). SNI increased the dynamic brush scores in both sexes on post-surgical days 3, 7, and 14, with females exhibiting lower scores than males (p<0.0001). These results confirm the development of mechanical hypersensitivity in PER2::LUC male and female mice following SNI, while showing sex-differences in the pain responses.

### Painful nerve injury decreases wakefulness and increases NREM sleep during the light phase in male PER2::LUC mice

To determine whether painful nerve injury alters sleep architecture in PER2::LUC male and female mice, we recorded longitudinal rodent polysomnography (rPSG) (Fig. 2A). The amount of time spent in each vigilance state was compared between baseline and day 14 within each group, and between the SNI and sham groups at day 14. To characterize baseline sex differences, epoch counts were also compared between male and female mice.

**Figure 2.**
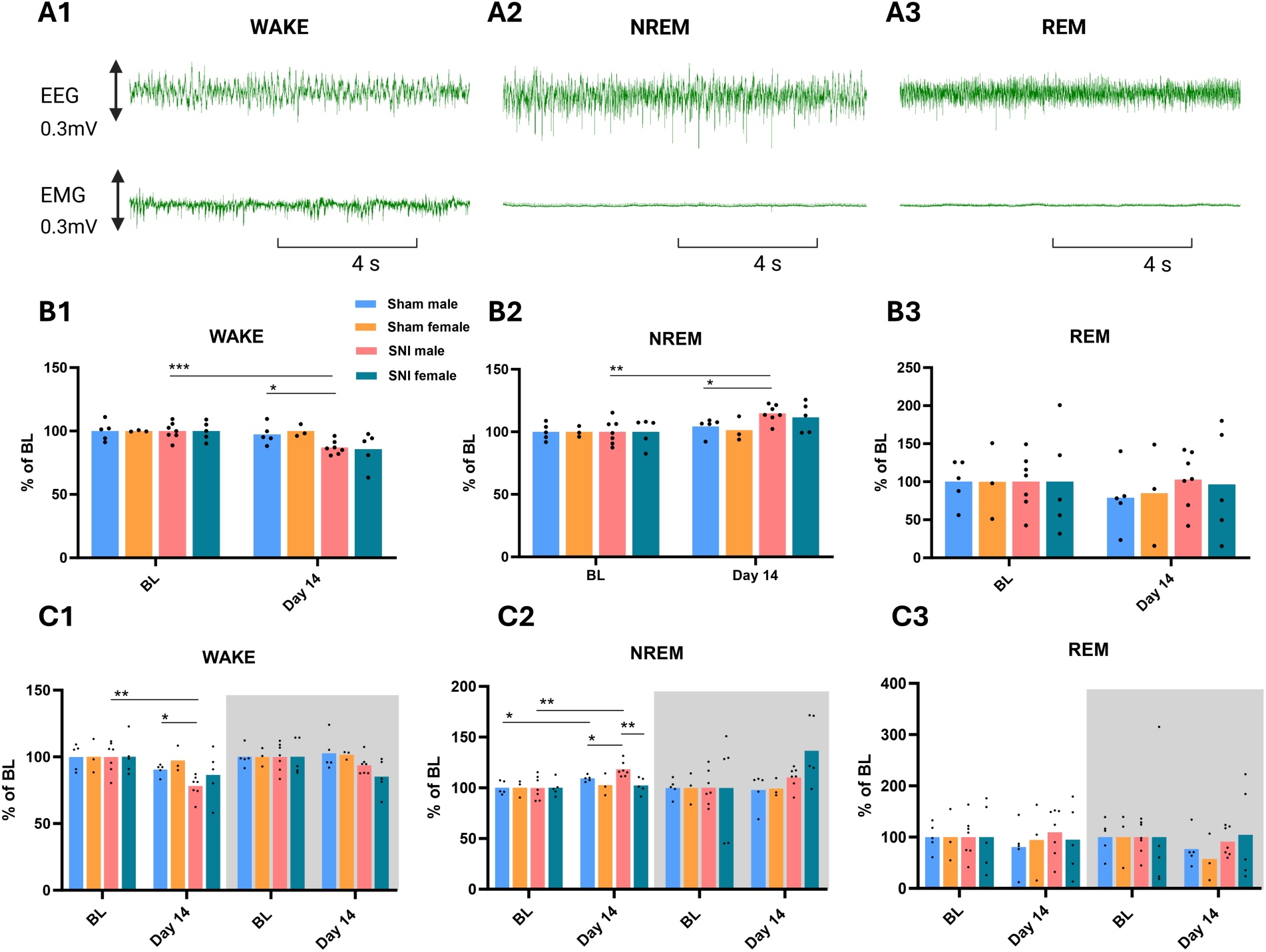
Distribution of sleep stages following spared nerve injury (SNI) or sham surgery in PER2::LUC male and female mice. **A)** Representative EEG and EMG recordings during wake (A1), non-rapid eye movement (NREM) sleep (A2), and rapid eye movement (REM) sleep (A3). **B)** Sleep-wake epochs (4 s) quantified across 24-h in sham and spared-nerve injury (SNI) males and females. SNI produced sex-dependent changes in sleep architecture, with significant alterations in wakefulness and NREM sleep detected only in males (B1 and B2). No significant alterations in REM epoch counts were observed between groups (B2) **C)** Wake, NREM, REM sleep analysed separately for the light and dark phase. During the light phase, SNI shifted the vigilance balance toward NREM sleep in males (C1 and C2), but not in females. Instead, SNI tended to increase NREM sleep during the dark phase in females. REM sleep did not differ significantly between groups (C3). The data is expressed as a percentage of the corresponding baseline values. Baseline and day 14 values within each group were compared using a two-tailed paired Student’s t-test, while differences between the SNI and sham at day 14 were analysed using a two-tailed unpaired Student’s t-test or Mann–Whitney test, depending on data normality. Individual dots represent measurements from individual animals. BL=baseline. *p<0.05, **p<0.01, ***p<0.001.

Female mice exhibited less non-rapid eye movement (NREM) sleep (t(18) = 3.7, p = 0.002) and more wakefulness (t(18) = 4.0, p = 0.0008) than male mice at baseline (Supplementary Fig. 1 A1 and A2). Separate analyses of the light and dark phases showed that these alterations occur during the active dark phase (wake: t(18) = 4.8, p = 0.0002; NREM: t(18) = 4.5, p = 0.0003) (Supplementary Fig. 1 B1 and B2). Additionally, while the total REM epochs remained unaffected (Supplementary Fig. 1 A3), females showed less REM sleep during the dark phase compared to males (t(18) = 3.0, p = 0.008) (Supplementary Fig. 1 B3).

SNI decreased wakefulness (t(6)=6.9, p=0.0004) and increased NREM sleep (t(6)=5.7, p=0.0012) in male mice at day 14 compared with baseline (Fig. 2 B1 and B2). Correspondingly, male SNI mice exhibited fewer wake epochs (t(10)=2.7, p=0.02) and more NREM epochs (U=4, p=0.03) than sham-operated mice at day 14, indicating that painful nerve injury shifts the vigilance state balance toward NREM sleep in PER2::LUC males.

Light- and dark-phase specific analyses revealed that these alterations were confined to the light phase, the resting phase in nocturnal animals (Fig. 2C). Longitudinal analyses of the light phase showed a reduction in wake (t(6) = 4.2, p=0.006) (Fig. 2 C1) and an increase in NREM sleep (t(6) = 4.01, p = 0.007) in male mice after nerve injury compared with baseline, as well as increase in NREM sleep in sham males (t(4)=2.8, p=0.049) (Fig. 2 C2). In the cross-sectional analyses at day 14, male SNI mice exhibited significantly more NREM sleep than male sham mice (t(10) = 2.5, p=0.031) and female SNI mice (t(10) = 3.6, p=0.005) during the light phase.

SNI did not significantly alter REM sleep in either sex (Fig. 2 B3 and C3). For wake and NREM sleep, however, no significant SNI-induced changes were detected in females (Fig. 2 B1-2, C1-2). Because SNI-induced changes in sleep-wake architecture were detected only in males, subsequent pharmacological experiments were performed in male mice.

**Figure 3.**
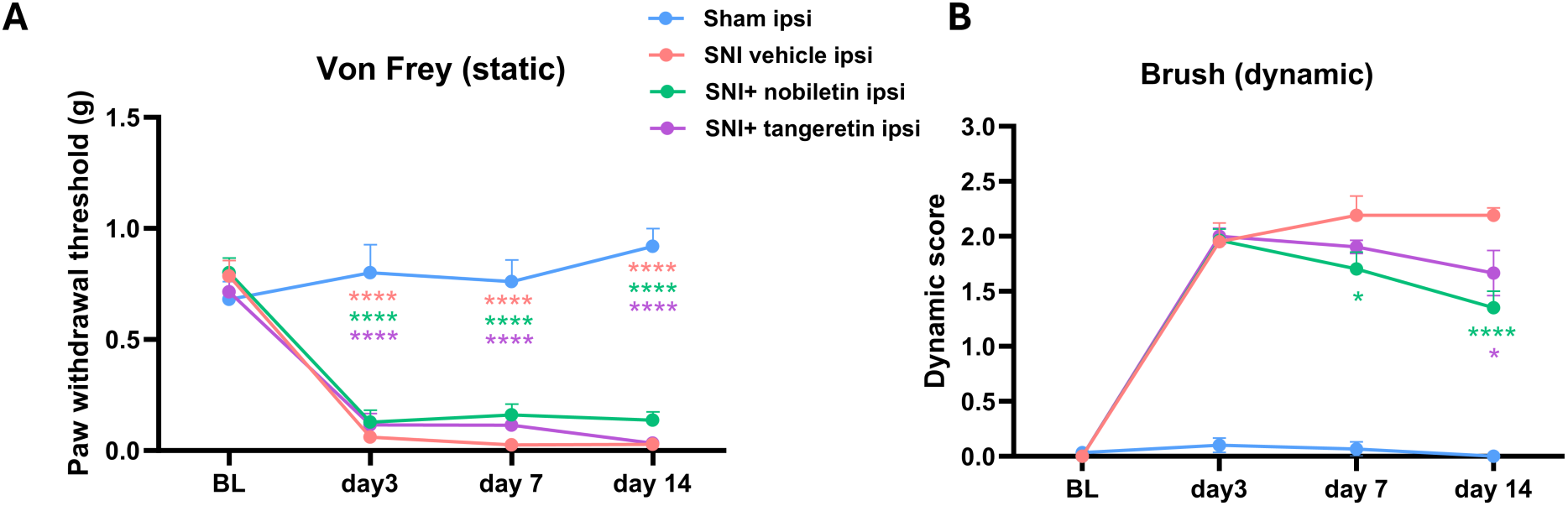
Impact of citrus-derived flavonoids nobiletin and tangeretin on pain thresholds at baseline and after sham or spared nerve injury (SNI) in male PER2::LUC mice. **A)** Von Frey results showed a significant post-surgical decrease in paw withdrawal thresholds in all groups. **B)** Nobiletin mitigated the effect of SNI on dynamic scores on day 7 and both flavonoids attenuated the effect on day 14. In the von Frey test, red asterisks indicate comparisons between the sham ipsi and SNI vehicle ipsi groups, green asterisks between the sham ipsi and SNI+ nobiletin ipsi groups, and purple asterisks between the sham ipsi and SNI+ tangeretin ipsi groups. In the brush test, green asterisks indicate comparisons between the SNI vehicle ipsi and SNI+ nobiletin ipsi groups, whereas purple asterisks indicate comparisons between the SNI vehicle ipsi and SNI+ tangeretin ipsi groups. Data were analysed using two-way ANOVA with Tukey’s post hoc test. The operated leg is referred to as ipsi (ipsilateral). BL = baseline. The sample sizes were n=7 for the SNI vehicle ipsi and the SNI+ tangeretin ipsi groups, n=5 for the sham ipsi group, and n=9 for the SNI+ nobiletin ipsi group. Mean ± SEM. *p<0.05, ****p<0.0001.

### Nobiletin and tangeretin alleviate injury-induced dynamic hypersensitivity in PER2::LUC male mice

To assess the impact of the citrus-derived flavonoids on pain-like behaviour upon SNI, osmotic minipumps delivering vehicle (10% DMSO in 90% saline), nobiletin (2 mg/kg/day), or tangeretin (1.8 mg/kg/day) were implanted immediately after SNI surgery in mice equipped with telemetric devices (Fig. 1A). Von Frey filament and brush tests were performed at baseline and at post-surgical days 3, 7, and 14.

Behavioural responses differed significantly over time between the experimental groups (von Frey (F(9, 96)=10.7, p<0.0001) and brush test (F(9, 96) = 14.75, p < 0.0001; Fig. 3A and B) (Fig 3 A and B). Nobiletin attenuated SNI-induced dynamic hypersensitivity at day 7 (p=0.01) and 14 (p<0.0001) and tangeretin mitigated dynamic scores at day 14 compared with SNI vehicle mice (p=0.01). Neither nobiletin nor tangeretin significantly altered paw withdrawal thresholds. The behavioural findings suggest that both flavonoids attenuate brush-evoked dynamic hypersensitivity following painful nerve injury.

### Tangeretin administration normalizes wake and NREM sleep following SNI

After establishing the SNI-induced changes in the sleep-wake cycle in male PER2::LUC mice with NP, we next evaluated whether flavonoid administration could reverse the abnormalities in the sleep architecture.

Within-group data comparisons between baseline and post-surgical day 14 showed a decrease in the wake epochs in the nobiletin group (t(8)=3.7, p=0.006), similarly to SNI vehicle animals, whereas tangeretin prevented this reduction (Fig. 4 A1). Consistently, wakefulness was lower in the SNI vehicle group than in tangeretin-treated SNI animals (t(12) = 3.2, p=0.008) at day 14. In addition, tangeretin-treated SNI mice had more wake epochs than nobiletin-treated SNI mice (t(14) = 2.2, P = 0.047). From baseline to day 14, NREM epochs increased in nobiletin-treated SNI animals (t(16) = 2.8, p=0.013), but not in SNI mice receiving tangeretin (Fig. 4 A2). At day 14, mice with nerve injury exhibited more NREM epochs than tangeretin-treated SNI mice t(7.9)= 3.8, p=0.005), with higher NREM epochs also observed in the nobiletin-treated SNI group compared to the tangeretin SNI mice (t(11.03)=2.3, p=0.04).

**Figure 4.**
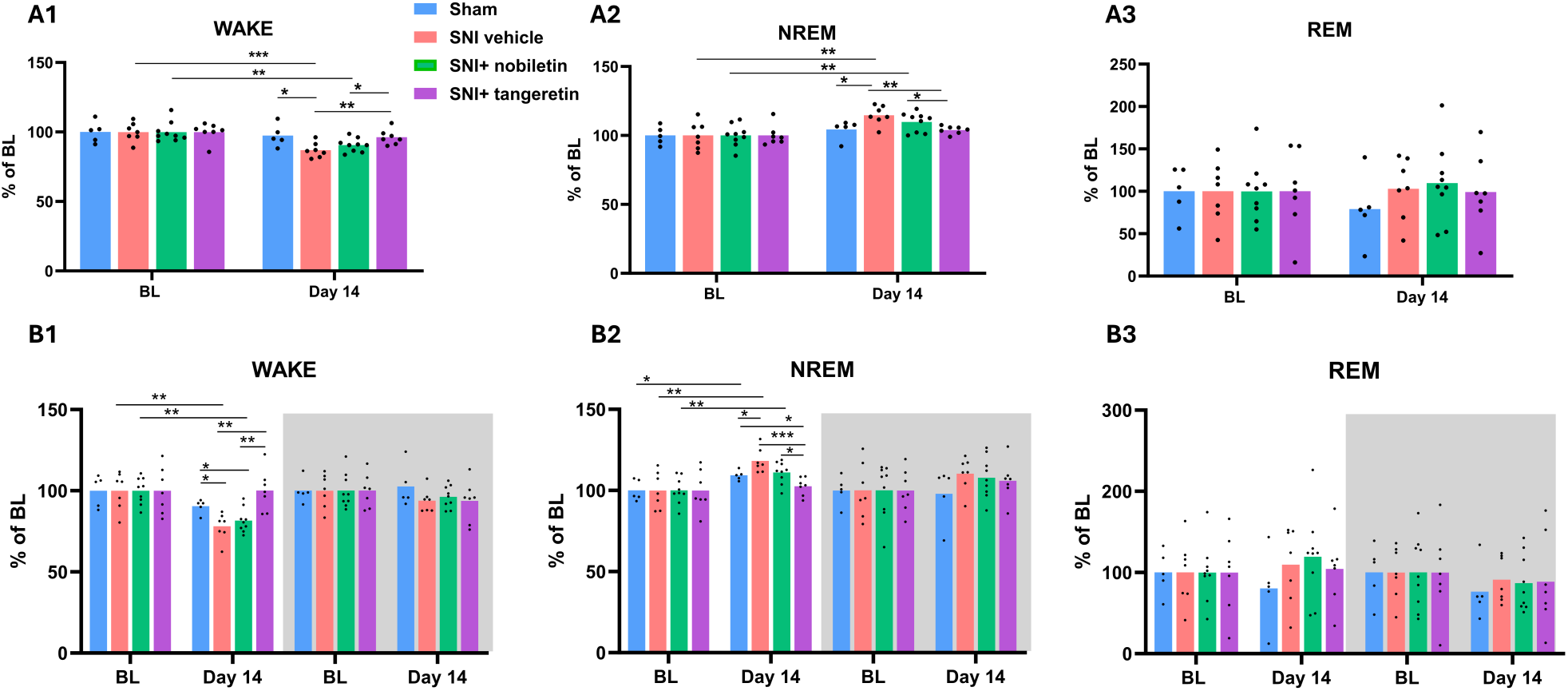
Effects of nobiletin and tangeretin on vigilance states in PER2::LUC males with painful nerve injury. **A)** Wake (A1), non-rapid eye movement (NREM) (A2), and rapid eye movement (REM) (A3) at baseline (BL) and post-surgical day 14 upon spared nerve injury (SNI) surgery across groups. Nobiletin and tangeretin differentially affected sleep-wake cycle following SNI, with tangeretin showing more pronounced effects. **B)** Vigilance states analysed separately during the light and dark phases (B1, B2, and B3). Flavonoid-induced alterations in the sleep-wake cycle were restricted to the light phase. Tangeretin reversed SNI-associated changes in wakefulness and NREM sleep. Differences in REM sleep between groups remained non-significant. The data are expressed as a percentage of the corresponding baseline values. Longitudinal comparisons were performed using paired t-tests, while group differences at day 14 were analysed using unpaired t-tests. Individual dots represent measurements from individual animals. *p<0.05, **p<0.01, ***p<0.001.

Light-dark phase analysis localized the alterations in wake and NREM to the light phase, with no detectable changes during the dark phase (Fig. 4 B1). Wake epochs declined from baseline to post-SNI day 14 in nobiletin-treated SNI animals (t(8) = 4.2, p=0.003) but not in tangeretin-treated mice. In line with this, nobiletin SNI group exhibited less wake epochs compared with sham (t(12)=2.4, p=0.03) and tangeretin-treated mice (t(14) = 3.63, p=0.003). Conversely, NREM epochs were higher in the nobiletin SNI group at day 14 than in the baseline (t(8)=4.8, p=0.001), as also observed in the SNI vehicle (Fig. 4 B2). At day 14, the tangeretin-treated mice exhibited fewer NREM epochs than the sham (t(10)=2.5, p=0.03), SNI (t(12)=4.5, p=0.0008) and nobiletin (t(14)=2.7, p=0.02) groups. Neither of the flavonoids significantly altered REM sleep (Fig. 4 A3 and B3). These findings indicate that tangeretin preserves normal sleep–wake architecture following nerve injury, whereas nobiletin does not.

### Tangeretin attenuates SNI-associated alterations in NREM spindle activity in male mice

To further investigate whether SNI-related alterations in NREM sleep extend to its microarchitecture, alterations in sleep activity, sleep spindles, short episodes of rhythmic EEG activity at sigma frequency, were quantified using automated algorithm^32^.

The results showed no significant changes in total spindle counts in the light phase, whereas during the dark phase nobiletin SNI group exhibited fewer spindles than SNI vehicle group (t(14) = 3.0, p=0.01) (Fig. 5A). SNI vehicle (t(6) = 8.07, p=0.0002) and nobiletin-treated SNI mice (t(8) = 3.41, p = 0.009) showed more NREM spindles at post-surgical day 14 than at baseline, whereas tangeretin administration attenuated this increase (Fig. 5C). Furthermore, the tangeretin SNI group had fewer NREM spindles compared with SNI vehicle (t(12)=2.8, p=0.017) and nobiletin SNI group (t(14)=3.3, p=0.005) at day 14.

**Figure 5.**
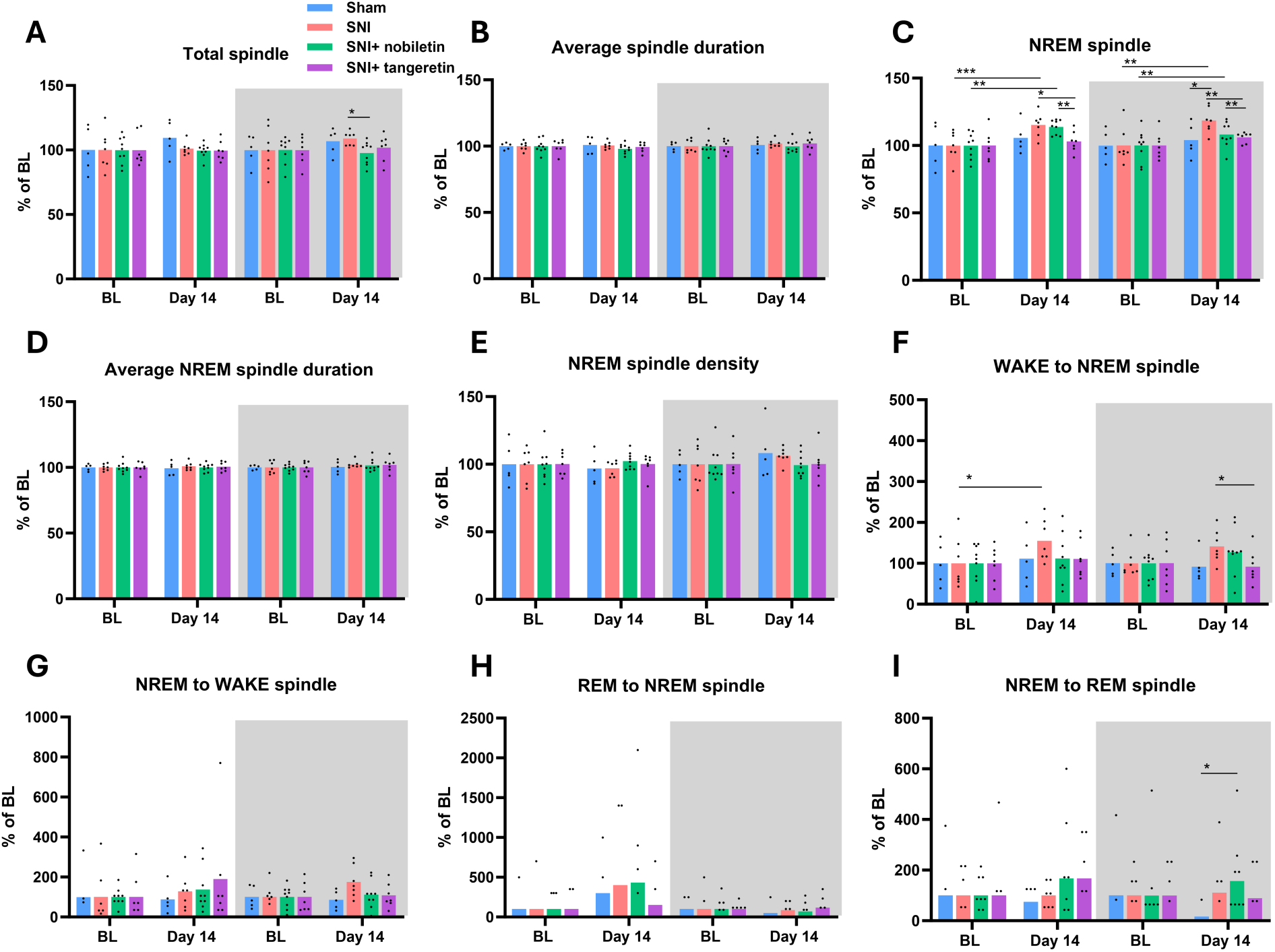
Changes in spindle activity upon SNI and flavonoid administration in PER2::LUC male mice. Light/dark phase-specific analysis across the 24-h recording period of **A)** total spindles, **B)** total spindle duration, **C)** non-rapid eye movement (NREM) spindles, **D)** NREM spindle duration E) NREM spindle density, calculated as the number of spindles per minute of NREM sleep, and transitions from **F)** wake to NREM, **G)** NREM to wake, **H)** REM to NREM, and **I)** NREM to REM. **A)** Nobiletin-treated SNI mice showed fewer total spindles than SNI vehicle mice during the dark phase, while no diffreences between groups were observed during the light phase. **C)** SNI increased NREM spindle activity during the light and dark phases, whereas tangeretin prevented this increase. **B, D)** Total and NREM spindle duration remained stable across experimental groups during both phases. **E)** No significant changes were observed in NREM spindle density between the groups **F)** SNI increased wake-to-NREM transitions from baseline to day 14 during the light phase, whereas tangeretin-treated SNI mice showed lower activity than SNI vehicle mice during the dark phase. **G, H)** NREM to wake and REM to NREM transitions remained unaffected by SNI or the flavonoids. **I)** During the dark phase, the nobiletin SNI group had more NREM to REM transitions than the sham group. BL= baseline. Data are shown relative to the corresponding baseline values, with baseline set to 100%. Paired t-tests were used for longitudinal analyses, whereas group comparisons at post-surgical day 14 were performed using unpaired t-tests or Mann–Whitney tests for non-parametric data. Individual dots represent measurements from individual animals. *p<0.05, **p<0.01, ***p<0.001.

The dark phase followed a similar pattern, with increased NREM spindles observed in within-group analyses of SNI vehicle (t(6) = 4.429, p = 0.004) and nobiletin-treated SNI mice (t(8)=3.4, p=0.009). At post-surgical day 14, the SNI vehicle group displayed more sleep spindles than the sham group (t(10)=2.3, p=0.04), while the tangeretin-treated SNI mice had fewer NREM spindles than SNI vehicle (t(12)=3.4, p=0.005) and nobiletin-treated SNI (t(14)=3.3, p=0.005) groups, demonstrating that tangeretin reversed the SNI-induced changes in NREM spindle numbers. However, NREM spindle density remained unchanged despite the increased number of NREM spindles (Fig. 5E). No significant changes were detected in the average total and NREM spindle durations (Fig. 5B and D).

Vigilance-state transitions were also impacted. SNI mice showed a significant increase in the proportion of wake to NREM transitions from baseline to day 14 during the light phase (t(6)=2.4, p=0.05) (Fig. 2F). In the dark phase, the longitudinal increase in the SNI vehicle group was not statistically significant (t(6)=2.052, p=0.086), while tangeretin-treated SNI mice showed fewer wake to NREM transitions than SNI vehicle mice at day 14 (t(12)=2.3, p=0.04). SNI tended to increase NREM to wake transitions during the active phase (Fig. 5G). At the light phase, NREM to REM and REM to NREM transitions were unaffected by SNI or the flavonoids (Fig. 5H and 5I). However, during the dark phase, nobiletin-treated SNI animals had a higher proportion of NREM to REM transitions (U = 7, p = 0.032).

### SNI increases EEG power across vigilance states, while flavonoids differentially mitigate these changes

Next, we assessed whether painful nerve injury alters the power spectra during wake, NREM, and REM sleep, and whether the flavonoids could reverse these changes and restore circadian dynamics in male mice. Power spectra of each vigilance state were analysed separately during the light (Fig. 6) and dark phases (Fig. 7). Alterations present at post-surgical day 14 but absent at baseline were described, along with longitudinal changes within groups across vigilance state-specific frequency ranges and 30-to 50-Hz low gamma range. All p-values and their respective degrees of freedom are presented in Supplementary table 1 for the light phase and Supplementary table 2 for the dark phase.

**Figure 6.**
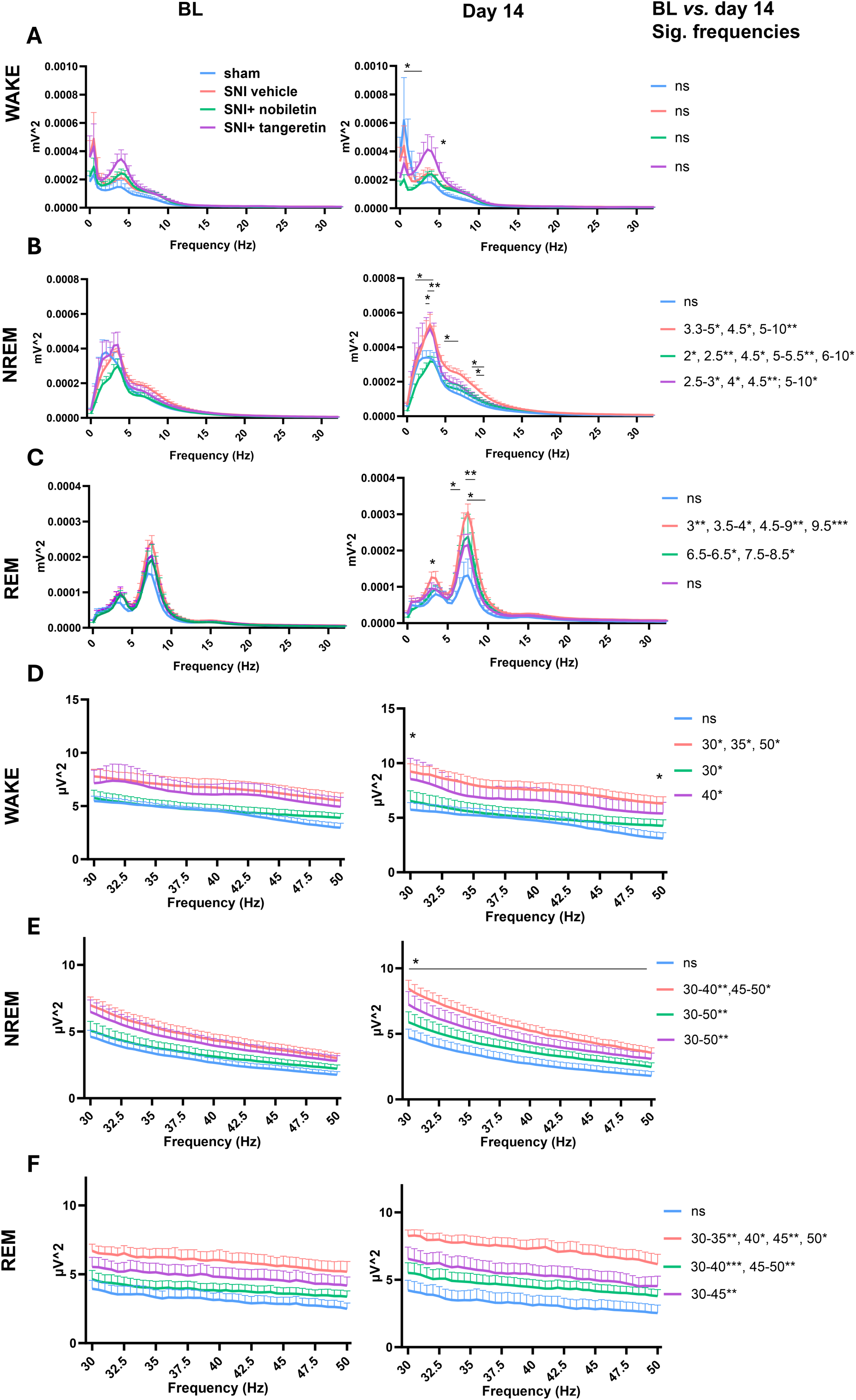
Changes in EEG spectral power during the light phase across vigilance states in PER2::LUC male mice. Spectral power was analysed across the 0.5–34.5 Hz frequency range **(A-C)** and the 30–50 Hz low-gamma range **(D-F)** at baseline and post-surgical day 14. Changes observed at post-surgical day 14 but not at baseline, as well as within-group changes between baseline and day 14, are reported. SNI increased EEG power mostly during NREM and REM sleep. During NREM sleep, nobiletin attenuated SNI-associated power increases around the dominant spectral peak, whereas tangeretin reduced power at higher frequencies. During REM sleep, both flavonoids attenuated SNI- associated increases, with tangeretin showing effects across a broader frequency range. Paired and unpaired t-tests were used for longitudinal and cross-sectional comparisons of power density, respectively. n=7 for the SNI vehicle and the SNI+ tangeretin groups, n=5 for the sham group, and n=9 for the SNI+ nobiletin group. BL= baseline. Mean ± SEM. *p<0.05, **p<0.01, ***p<0.001.

**Figure 7.**
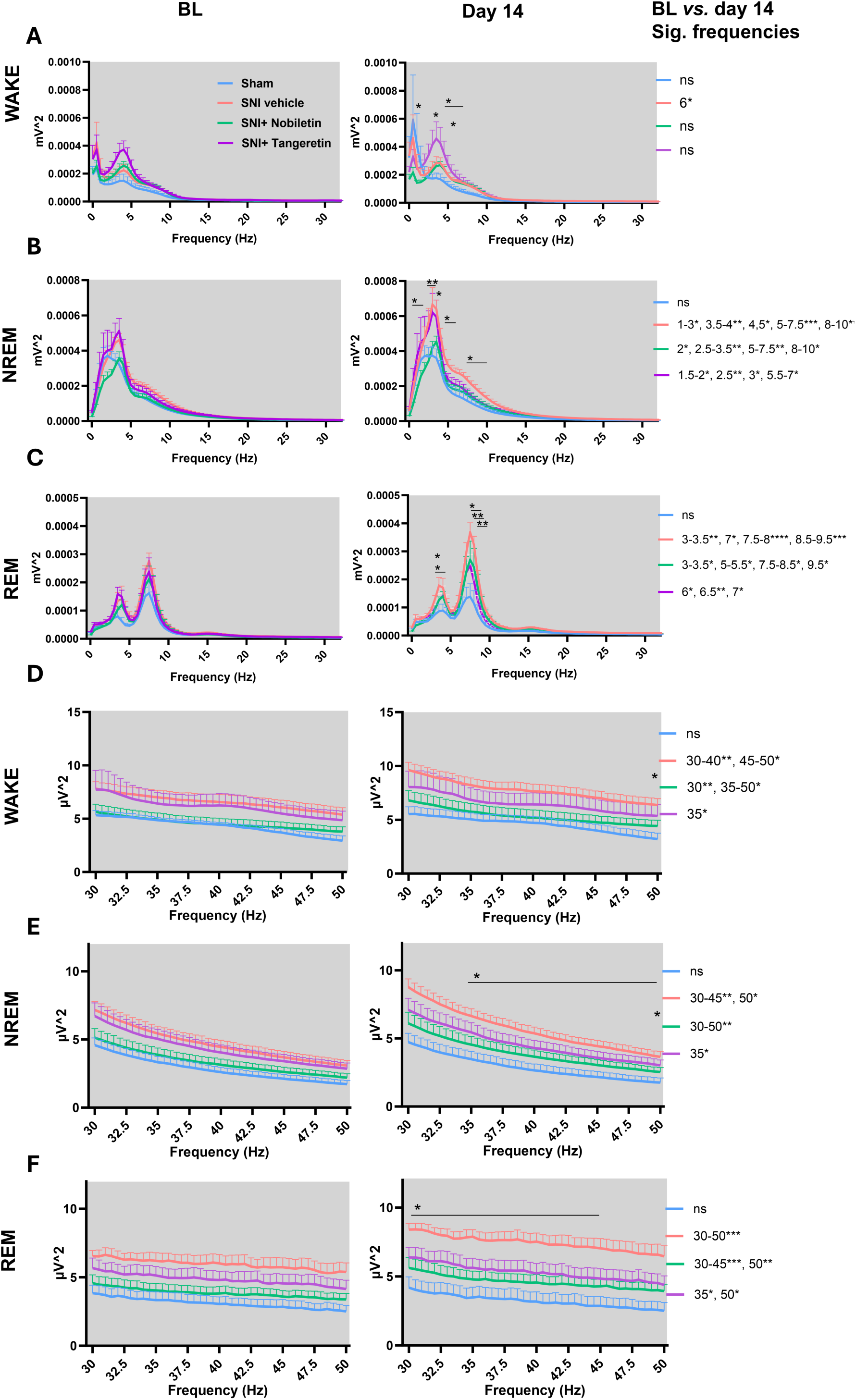
Changes in EEG spectral power during the dark phase across vigilance states in PER2::Luc male mice. Power spectra were examined during wake, non-rapid eye-movement (NREM), and rapid eye movement (REM) states at baseline and post-surgical day 14. Statistical comparisons included differences between experimental groups at day 14 and longitudinal changes within each group from baseline to day 14. During the dark phase, SNI increased EEG power across all vigilance states in the 0.5–34.5 Hz frequency range (A–C), and in the 30–50 Hz low-gamma range (D–F). Nobiletin and tangeretin administration attenuated SNI-associated EEG power increases in a frequency-specific manner, with tangeretin affecting a wider range of frequencies than nobiletin. Changes in power density over time were tested with paired t-tests, whereas unpaired t-tests were applied to day 14 group comparisons. n=7 for the SNI vehicle and the SNI+ tangeretin groups, n=5 for the sham group, and n=9 for the SNI+ nobiletin group. BL= baseline. Mean ± SEM. *p<0.05, **p<0.01, ***p<0.001.

### Power spectral analysis during wake

EEG power spectra was altered following SNI in all vigilance states, whereas after sham EEG power remained stable across frequencies and vigilance states throughout the study. SNI surgery and the flavonoid administration did not alter wake EEG power longitudinally, while SNI vehicle exhibited higher EEG power at 6 Hz compared with sham mice, and tangeretin SNI mice showed higher power than nobiletin SNI mice in the low delta1.0-2.5 Hz range at day 14 during the light phase (Fig. 6A). Nerve injury increased the low gamma power at 30-35 Hz and 50 Hz, while increases in the nobiletin and tangeretin-treated SNI groups were detected only at 30 and 40 Hz, respectively (Fig. 6D). At day 14, nobiletin SNI mice showed lower power than SNI vehicle mice at 30 and 50 Hz.

During the dark phase, wake 6-Hz activity was elevated in SNI vehicle mice at post-surgical day 14 compared with baseline (Fig. 7A). At day 14, relative to sham, SNI vehicle mice showed increased power across 5-6 Hz, while nobiletin-treated SNI at 4 and 6 Hz. In addition, tangeretin-treated SNI mice exceeded nobiletin SNI mice at 1 Hz. For low-gamma power, SNI induced a broad increase across frequencies (30-50 Hz), which was not reversed by nobiletin (30-50 Hz) but was absent following tangeretin treatment, except at 35 Hz (Fig. 7D). At day 14, SNI vehicle mice showed higher power than nobiletin-treated SNI mice at 50 Hz. Overall, SNI was associated with increased wake EEG power, and these alterations were largely mitigated by tangeretin administration.

### Power spectral analysis during NREM

SNI vehicle, nobiletin SNI and tangeretin SNI groups displayed longitudinal increases in NREM power (SNI: 3–3.5, 4.5–10 Hz; nobiletin: 2–2.5, 4.5–10 Hz; tangeretin: 2.5–3, 4–10 Hz) and across the entire low-gamma range (30–50 Hz) during the light phase (Fig. 6B and E). By day 14, nobiletin SNI mice showed reduced power relative to SNI vehicle mice at 0.5–2.0, 3.5–4, and 9.5-10 Hz, including the region of the prominent spectral peak, whereas tangeretin SNI group had lower power across the higher 8.5-10 Hz range. SNI vehicle mice had greater NREM power than sham mice at 3.0-6.5 Hz and higher low-gamma power than nobiletin-treated SNI mice throughout the 30-50 Hz range.

During the dark phase, longitudinal increases in NREM power extended across 1–10 Hz in SNI vehicle mice and 2.0-3.5 and 5-10 Hz in nobiletin-treated SNI mice, whereas in the tangeretin-treated SNI mice changes occurred at 1.5–3 and 5.5–7 Hz (Fig. 6B). At post-surgical day 14, the differences across frequencies paralleled those observed during the light phase. SNI vehicle mice exceeded sham at 3 and 5-6 Hz, whereas NREM power was lower in nobiletin SNI mice at 0.5–1.5 and 3.5–4 Hz and in SNI tangeretin mice at 7.5–10 Hz than in SNI mice. Low-gamma power also increased longitudinally across 30–50 Hz in SNI vehicle and nobiletin SNI mice, with tangeretin-treated SNI showing a change only at 35 Hz (Fig. 7E). At day 14, tangeretin SNI group had higher low-gamma power than sham group at 50 Hz, while SNI vehicle mice exceeded nobiletin SNI mice across 35–50 Hz. These results indicate that nobiletin attenuated the SNI-induced NREM power increase around the dominant spectral peak, whereas tangeretin exerted its strongest effects at higher frequencies.

### Power spectral analysis during REM

REM power showed no changes from baseline to day 14 in tangeretin-treated SNI mice, whereas SNI vehicle (3-9.5 Hz) and nobiletin-treated SNI mice (6.0–6.5, and 7.5–8.5 Hz) showed elevated REM power across a range of frequencies during the light phase (Fig. 6C). Correspondingly, a postsurgical increase in REM EEG power was observed in SNI vehicle animals compared with sham (3.0 Hz, 5.5-7.0 Hz). SNI mice receiving flavonoids displayed lower EEG power than SNI vehicle mice, with differences observed at 3 Hz for nobiletin and 7.5–9.5 Hz for tangeretin administration. Low-gamma power increased longitudinally across 30–50 Hz in SNI vehicle and nobiletin-treated SNI mice and across 30–45 Hz in tangeretin-treated SNI mice (Fig. 6F).

During the dark phase, within-group increases in REM power occurred in SNI vehicle (3–3.5 and 7– 9.5 Hz), nobiletin SNI (3–3.5, 5–5.5, 7.5–8.5, and 9.5 Hz), and tangeretin SNI mice (6–7 Hz). At day 14, REM power was elevated in SNI vehicle mice relative to sham mice at 3–3.5 and 8–9 Hz, while nobiletin attenuated this elevation at 3 Hz and tangeretin attenuated it across 7.5–9.5 Hz frequency range (Fig. 7C). Postsurgical low-gamma power rose across 30–50 Hz in SNI vehicle and nobiletin-treated SNI mice but only at 35 and 50 Hz in tangereting-treated SNI mice (Fig. 7F). Furthermore, low gamma power was markedly higher in the SNI vehicle group than in the tangeretin SNI group (30- 45Hz). To conclude, SNI increased REM EEG power across both light and dark phases, while the flavonoids attenuated these changes in a frequency- and phase-dependent manner, with tangeretin most effectively attenuating the SNI-related power increases.

### Nobiletin and tangeretin restore Per2 amplitude in dorsal root ganglia to pre-surgical levels in male mice

To investigate whether SNI alters circadian *Per2* rhythmicity in pain-regulating DRGs and whether these changes in *Per2* rhythms can be reversed by flavonoid administration, lumbar ipsilateral DRGs were collected on post-surgical day 14 for bioluminescence recordings in LumiCycle 32 device (Actimetrics, Lafayette, IN, USA) and the changes in light intensity were monitored continuously for 5 days (Fig. 8A). Sham-operated mice exhibited a robust *Per2* circadian rhythmicity, characterized by average amplitude of 6.7 ± 2.3 counts/sec, period of 23.7 ± 0.1 h, and acrophase of 19.9 ± 0.7 h (Fig. 8B-E). One-way ANOVA revealed a significant difference in amplitude (F(3,12) = 6.9, p=0.006) and period (F(3, 12) = 3.898, p = 0.037) between groups (Fig. 8F and G). SNI surgery significantly increased the *Per2* amplitude in DRGs compared to sham operated mice (45,0 ± 12.4 counts/sec, p=0.02), indicating that circadian rhythmicity is dysregulated under NP conditions. Administration of nobiletin (5.2 ± 1.4 counts/ sec, p=0.015) and tangeretin (2.9 ± 0.8 counts/ sec, p=0.018) reversed the increased *Per2* amplitude, demonstrating their clock-resynchronizing effect in mice with NP.

**Figure 8.**
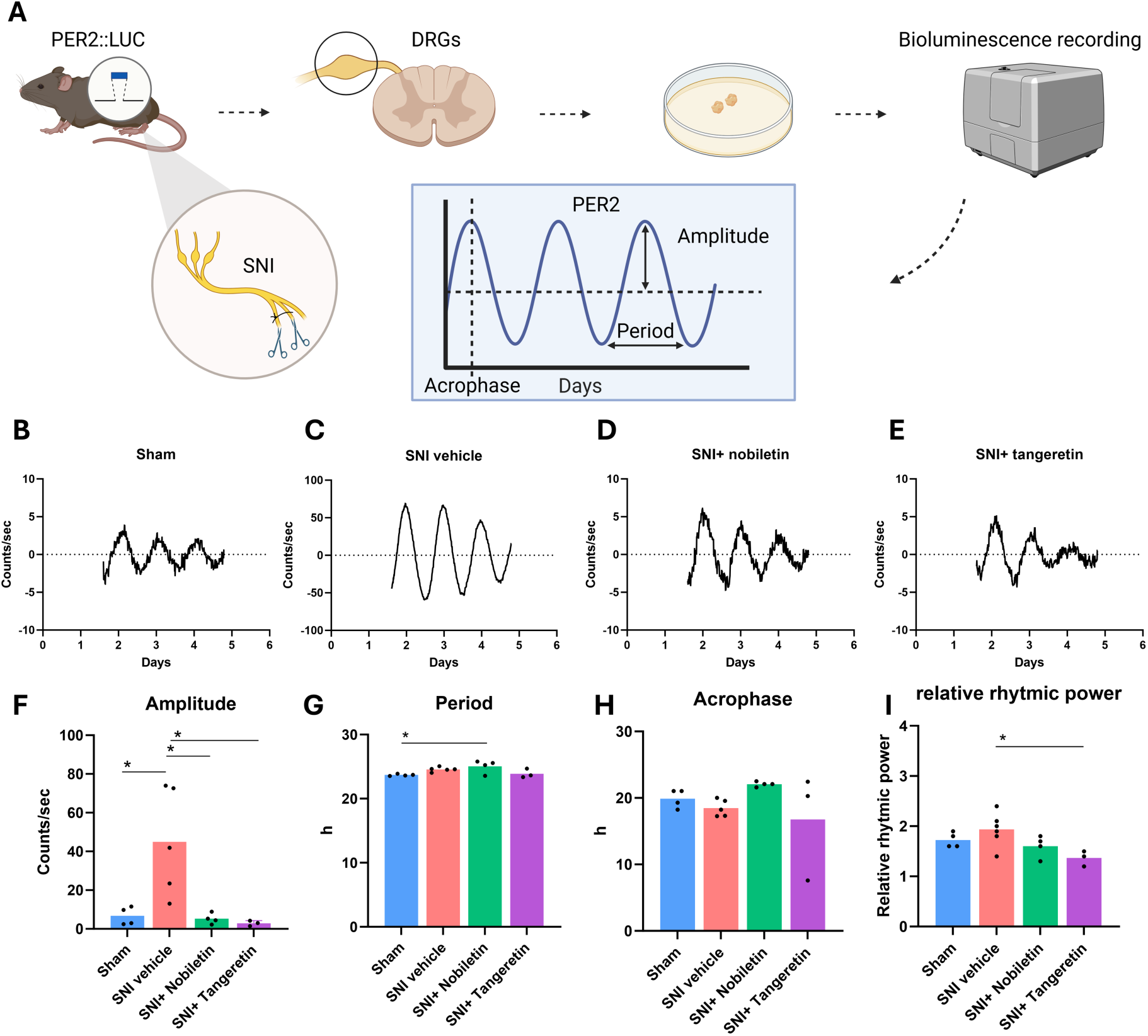
Effects of nerve injury on circadian oscillations in the ipsilateral dorsal root ganglia in PER2::LUC male mice. **A)** Experimental workflow and analysed circadian parameters. DRGs were collected following spared nerve injury (SNI) or control sham surgery and placed in the LumiCycle system for continuous bioluminescence recording over 5 days. A representative real-time bioluminescence recording is shown for each experimental group: **B)** sham, **C)** spared nerve injury, **D)** SNI+ Nobiletin and **E)** SNI+ Tangeretin. Circadian rhythm parameters **F)** amplitude (distance from midline to the peak), **G)** acrophase (timing of the peak), **H)** period (duration of one cycle), and **I)** relative rhythmic power (strength of rhythmicity) were quantified using LumiCycle Analysis software (version 3.103). **F)** SNI enhanced *Per2* amplitude in the DRGs versus sham-operated mice, whereas both flavonoid treatments restored the amplitude to pre-surgical levels. **G and I)** The nobiletin-treated SNI mice exhibited a longer period than sham mice, while the SNI vehicle group showed higher relative rhythmic power compared with tangeretin-treated SNI mice. **H)** No significant alterations were observed in the acrophase between groups. Circadian parameters are presented as mean ± SEM and were analysed by one-way ANOVA followed by Tukey’s post hoc test. *p<0.05. Figure A was created using BioRender.

A non-significant advance in acrophase was observed in SNI vehicle group (18.5 ± 0.6 h) compared to the sham group (19.9 ± 0.7 h) (Fig. 8H). Nobiletin tended to delay the acrophase (22.1 ± 0.2 h) and lengthened the period (25.0 ± 0.5, p=0.042) compared to sham mice. No significant differences were observed between sham, SNI vehicle (24.7 ± 0.1 h) and tangeretin-treated SNI mice (23.9 ± 6.9 h) in period length. Samples exhibited relative rhythmic power across all groups, with significant difference between SNI vehicle (1.9 ± 0.3) and tangeretin-treated SNI mice (1.4 ± 0.2, p=0.03) (Figure 8I).

## Discussion

With circadian disruption increasingly implicated in NP^2,13–16^, understanding these circadian abnormalities is essential for developing new chronotherapeutic approaches for pain relief. However, progress has been limited by experimental approaches that require tissue collection at multiple time points across the circadian cycle^13–15^, providing only snapshots of molecular clock activity rather than its continuous dynamics. To address this, we introduced SNI to circadian reporter PER2::LUC mice to establish a model that allows real-time monitoring of *Per2* rhythmicity in pain-regulating sensory tissues. In PER2::LUC males, this model captured disturbances in both DRG clock rhythmicity and the sleep-wake cycle under NP conditions, with flavonoid administration restoring *Per2* rhythmicity in DRGs to pre-surgical levels, reversing SNI-induced sleep changes, and attenuating dynamic hypersensitivity post-surgery.

SNI induced mechanical hypersensitivity in both sexes of PER2::LUC mice, confirming the development of NP phenotype. At baseline, the PER2::LUC model recapitulated sex differences in sleep-wake architecture previously reported in C57BL/6J mice^33,34^, with females exhibiting more wakefulness and less NREM sleep during the dark phase. To our knowledge, this is the first demonstration that SNI disrupts the sleep–wake cycle in freely moving PER2::LUC male mice. In females, no significant SNI-induced changes in sleep architecture were detected. This may reflect limited statistical power or sex-specific biological factors, such as hormones and estrous cycle-dependent variation in sleep-wake regulation^35^. Interestingly, SNI shifted the sex-specific pattern observed at baseline, with male mice exhibiting more NREM sleep than females, suggesting that NP alters sleep-wake regulation differently in male and female PER2::LUC mice. Because SNI-induced sleep changes were detected only in males, subsequent pharmacological experiments were performed in male mice to test whether these alterations could be reversed.

The effects of NP on sleep architecture appear to be model-dependent, with previous studies reporting reduced^20,36–38^ or unchanged^16,39^ NREM sleep, unchanged^39^ or increased wakefulness^20,36–38^, and reduced^16^, unchanged^20,36,38,39^ or phase-specific increases in REM sleep^37^. Interestingly, SNI increased NREM sleep and reduced wakefulness in PER2::LUC male mice, contrasting with the sleep–wake alterations reported in most previous studies. This pattern may reflect characteristics of the transgenic PER2::LUC model or represent a compensatory increase in NREM sleep in response to persistent pain. However, increased NREM sleep does not necessarily indicate restorative sleep. SNI increased wake-to-NREM transitions, while NREM-to-wake transitions showed a similar trend during the dark phase, suggesting altered sleep stability. This is in line with a recent study showing increased NREM sleep arousability in SNI mice^40^. Moreover, ongoing ectopic activity from injured peripheral nociceptors in mice with SNI drives instability in sleep microstructure, fragmenting sleep continuity and forcing an abnormal, fragmented sleep state, despite the increase in number of NREM sleep episodes^41^. Alternatively, increased NREM sleep may also be relevant to pain processing, as nerve injury has been shown to activate a neural pathway involving the parabrachial nucleus, basal forebrain, and somatosensory cortex during NREM sleep, promoting mechanical allodynia^42^. Thus, whether the increased NREM sleep represents a compensatory or pathological response to painful nerve injury remains unclear.

The circadian-modulating tangeretin restored SNI-induced alterations in sleep-wake architecture to presurgical levels, supporting its potential to stabilize sleep upon painful nerve injury. Moreover, tangeretin also reversed NP-induced changes in sleep-wake transitions, suggesting improved sleep stability. Prior to the present study, the effects of tangeretin on sleep have remained largely unexplored, with only a single study reporting sleep-promoting effects, including shortened sleep onset latency and extended sleep duration in a thiopental-induced sleep model in mice^28^. Although nobiletin did not restore the overall sleep-wake architecture, it reversed SNI-induced changes in wake to NREM transitions during the light phase, indicating differential beneficial effects of these closely related flavonoids on sleep under NP conditions.

SNI also enhanced EEG spectral power across vigilance states, showing that nerve injury does not only affect sleep architecture but also vigilance state-dependent brain activity^43^. SNI increased NREM spectral power across multiple frequencies, including the delta range (1–4 Hz). As NREM delta power is regarded as a marker of sleep intensity and homeostatic sleep pressure^44^, its increase following SNI may indicate an increased need for sleep during NP. Nobiletin attenuated this increase, whereas tangeretin exerted its strongest effects at higher frequencies. SNI increased wake EEG spectral power, particularly in the low-gamma range (30-50 Hz), with similar increases in low-gamma activity also observed during NREM and REM sleep. Gamma activity is associated with cortical arousal during wakefulness and sleep^45,46^, linked to pain perception and positively correlates with pain intensity in humans^47^. Thus, the widespread increase in gamma activity may reflect altered cortical processing following nerve injury. Notably, tangeretin broadly attenuated SNI-induced increases in low-gamma activity across vigilance states. SNI increased REM EEG power within the theta range during both light and dark phases, a phenomenon previously observed in both clinical and preclinical studies of NP and linked to altered activity within brain networks involved in pain processing^37,48^. Tangeretin largely prevented these SNI-induced alterations, particularly during the light phase.

Sleep spindles are short periods of thalamocortical activity during NREM sleep that support sensory processing, neural plasticity, and memory formation^49^. Spindle activity increased during NREM sleep following SNI, whereas this increase was absent in tangeretin-treated SNI mice. However, as spindle density remained unchanged, the enhanced spindle activity most likely reflects the increased duration of NREM sleep rather than increased spindle formation. Overall, these findings show that SNI disturbs multiple aspects of sleep-wake cycle, with nobiletin attenuating alterations in sleep-wake transitions and NREM spectral activity and tangeretin exerting wider sleep-restoring effects, including normalization of the sleep architecture.

Both nobiletin and tangeretin attenuated SNI-induced dynamic hypersensitivity in PER2::LUC males. This extends the previously reported antinociceptive effects^26,27^ of nobiletin to the SNI model of NP and to brush-evoked dynamic mechanical hypersensitivity, while providing evidence that tangeretin can similarly modulate NP-induced sensory abnormalities. In contrast, paw withdrawal thresholds in von Frey filament tests remained unaffected in the PER2::LUC SNI mice upon flavonoid administration. Dynamic mechanical allodynia is primarily mediated by Aβ low-threshold mechanoreceptors (LTMR), whereas localized punctate mechanical sensitivity involves Aβ, Aδ, and C afferents^50^. Given that these two forms of mechanical hypersensitivity rely on partially distinct fibres, the attenuation of dynamic hypersensitivity suggests that the flavonoids may affect the processing of Aβ-LTMR signalling rather than inducing general mechanical analgesia. Consistent with this interpretation, dynamic and punctate mechanical hypersensitivity can occur independently in patients with peripheral NP and show differential responses to sensory nerve blockade supporting their distinct underlying mechanisms^51^. Such sensory differences are also evident in postherpetic neuralgia, where dynamic mechanical hypersensitivity and altered pinprick sensitivity differ between patient subgroups and may change independently over time^52^. Alternatively, the different effects on dynamic and punctate mechanical hypersensitivity may be related to differences in dose, pain model, route of administration, or time of testing. To avoid disturbing circadian rhythmicity and sleep-wake cycle, we administered nobiletin continuously using osmotic minipumps, whereas a 60-fold higher dose of nobiletin, administered to mice by oral gavage, previously attenuated von Frey paw withdrawal responses in a CCI model^26^. In addition, pain sensitivity varies across the day^31^, and analgesic effects may therefore depend on the time of testing. Given that SNI is associated with neuroinflammation in pain-regulating tissues ^53^, the anti-inflammatory properties of nobiletin and tangeretin^54^ may also have contributed to the reduction in dynamic mechanical hypersensitivity.

In the PER2::LUC SNI model, DRGs exhibit robust *Per2* rhythmicity, further supporting the presence of a local peripheral circadian clock in the pain-regulating DRGs^55^. Notably, SNI surgery increased the amplitude of *Per2* rhythms in DRG explants, providing evidence that NP is associated with altered circadian clock dynamics. Recent data demonstrates that the molecular clock within DRG nociceptors regulates neuronal excitability and circadian variation in nociceptive sensitivity^31^. Thus, disturbances in local clock function could directly affect nociceptive processing. Our findings further support this possibility, as both nobiletin and tangeretin prevented the SNI-induced increase in amplitude of *Per2* rhythms in DRGs while attenuating dynamic hypersensitivity. Given that only tangeretin restored the sleep-wake architecture, these findings point to altered DRG clock dynamics as a potential contributor to the sensory phenotype rather than a consequence of restored sleep-wake organization. Since disturbed *Per1* rhythmicity has been reported in the spinal dorsal horn of mice subjected to partial sciatic nerve ligation^15^, the flavonoid-induced clock resynchronization may not be limited to DRGs but could also occur in other regions involved in pain processing, such as the spinal dorsal horn. Nobiletin and tangeretin have previously shown strong molecular clock-modulating effects^23–25,56^. For instance, nobiletin improved disrupted clock-gene rhythms in the liver of mice fed with a high-fat diet^23^, potentially via retinoic acid receptor-related orphan receptor signalling, while both nobiletin and tangeretin increased PER2::LUC amplitude in mouse embryonic fibroblasts^24^. In a mouse model of Alzheimer’s disease, nobiletin altered cortical clock gene expression, but did not significantly improve sleep^57,58^, suggesting that modulation of the circadian clock may not necessarily translate into changes in vigilance states. However, in the present study, nobiletin also attenuated the SNI-induced increase in NREM EEG power, indicating that its effects were not restricted to peripheral clock rhythmicity but extend to sleep-related brain activity.

The therapeutic potential of circadian biology remains largely underexplored in pain research^5^. Evidence of this potential is already apparent in rheumatoid arthritis, where aligning glucocorticoid treatment with the disturbed diurnal rhythm of tumour necrosis factor-α (TNF-α) markedly improved clinical outcomes and reduced joint pain^59,60^. Moreover, non-pharmacological chronotherapies such as light therapy, which directly regulates the circadian master clock^61^, have shown beneficial effects in painful conditions^62,63^ and pain-related comorbidities such as sleep disturbances^61^. Sleep and pain interact bidirectionally, and improving sleep may also improve pain outcomes^2^. Some pharmacological treatments for NP, such as pregabalin, may improve sleep^2,16^, whereas others, including certain antidepressants, may contribute to sleep disturbances^2^. Considering the beneficial effects of the flavonoids on circadian regulation and dynamic mechanical hypersensitivity, co-administration of these compounds with analgesic treatments that may adversely affect sleep could allow for dose-reduction of standard drugs, potentially limit treatment-related side effects and restore the sleep-wake cycle. More broadly, the involvement of multiple circadian processes in NP, including sleep regulation and molecular clock function, further expands the potential for chronotherapeutic interventions but also introduces complexity. Targeting these abnormalities may improve disease outcomes, while their complex interplay makes it challenging to determine how individual processes contribute to therapeutic responses. Defining these relationships may help to guide the development of chronotherapeutic strategies for NP.

Our study has some limitations. The absence of a significant SNI-induced sleep phenotype in females should not be interpreted as evidence of no effect due to the reduced final sample size in sham group and potential effects of estrous cycle variation and gonadal hormones. In addition, sleep was assessed on post-surgical day 14, and sex-specific changes in sleep architecture may emerge earlier or later under NP conditions. Moreover, the pharmacological and *Per2* bioluminescence experiments were restricted to males based on the male-specific changes upon SNI, so whether nobiletin and tangeretin are effective in females remains unknown. Due to solubility properties of flavonoids and dosage limitations of using the osmotic minipumps, only the effects of a single dose of each flavonoid were tested. Finally, the PER2::LUC reporter is a knock-in fusion model, and its effects on sleep and pain behaviour relative to wild-type mice were not assessed. Although a comprehensive wild-type comparison has not been reported, NREM sleep duration and EEG delta power dynamics in PER2::LUC mice appear to be similar to those in C57BL/6J mice, which share much of the PER2::LUC genetic background^64^. Larger studies, incorporating estrous cycle monitoring, additional time points, and both sexes, will be needed to assess the generality of these findings.

Overall, PER2::LUC SNI mouse model overcomes the limitations of snapshot-based approaches requiring tissue collection across multiple circadian time points by providing a means to investigate how peripheral clock dynamics evolve under NP conditions and respond to chronobiotic interventions in real time. We showed that SNI disturbs *Per2* rhythmicity in the DRGs and sleep–wake cycle, whereas flavonoid chronotherapy counteracts these alterations and attenuates dynamic hypersensitivity. These findings support a circadian basis of NP, demonstrate that SNI-induced disturbances in *Per2* rhythmicity are chronopharmacologically targetable and highlight the chronotherapeutic potential of nobiletin and tangeretin in NP.

## Material and methods

### Animals

The experiments were performed on 38 homozygous mutant PER2::LUC mice (28 males and 10 females, 15 ± 0.5 weeks of age) obtained from The Jackson Laboratory (strain 006852: Bar Harbor, ME, USA). The PER2::Luc mouse model, with fusion of circadian *Per2* with luciferase, allows for real-time bioluminescence monitoring of circadian changes in the tissues of interest *ex vivo*^30^. The colony was established at the Laboratory Animal Centre at the Viikki Campus of the University of Helsinki. Animals were individually housed to ensure reliable EEG/EMG recordings under standard 12:12h light/dark cycle. Cage locations within the room were randomized and changed weekly to minimize potential environmental effects. The animals had *ad libitum* access to standard rodent chow (Teklad 2916 rodent chow, Envigo, United Kingdom) and water throughout the study. All animal procedures were approved by the Regional State Administrative Agency for Southern Finland (ESAVI/4935/2024) and study complied with the ARRIVE guidelines and the EU Directive 2010/63/EU for animal experiments.

### Genotyping

Prior to the experiments, all mice were genotyped to confirm homozygosity by PCR using ear punches at the Genetically Modified Rodents (GM) Unit, University of Helsinki. The DNA lysate was prepared using the Extracta DNA Prep for PCR kit (QUNT95091-002, Quantabio) according to the manufacturer’s instructions, and PCR amplification was performed using DreamTaq DNA Polymerase (EP0705, Thermo Fisher) in the SimpliAmp™ Thermal Cycler (A24811, Thermo Fisher). Two uL of lysate were used for step-down PCR: 95°C for 3 min; 5 cycles of 95°C for 30s, 66°C for 25 s, and 72°C for 35 s; 5 cycles of 95°C for 30s, 64°C for 25 s, and 72°C for 35 s; 25 cycles of 95°C for 30, 62°C for 25 s, and 72°C for 35 s. The strain-specific genotyping primer sequences for PER2::LUC were based on Jackson Laboratory data. To identify homozygotes, the common forward (oIMR6588)-5’ CTGTGTTTACTGCGAGAGT and mutant reverse (oIMR6590)-5’ TAAAACCGGGAGGTAGATGAGA were applied. For the wild-type allele, the common forward (oIMR6588)-5’ CTGTGTTTACTGCGAGAGT and wild-type reverse (oIMR6589)-5’ GGGTCCATGTGATTAGAAAC were applied. The PCR products were separated on a 1.5% agarose gel to distinguish the PER2::LUC genotypes, with the wild-type band at 230 bp, the mutant at 650 bp, and the heterozygous showing both 230 bp and 650 bp bands.

### Study protocols

Male (SNI n=7, sham n=5) and female (SNI n=5, sham n=5) PER2::LUC mice were randomly assigned to the experimental groups and implanted with EEG/EMG transmitters two weeks before SNI or control sham surgeries. SNI males continuously received vehicle (n=7), nobiletin (2 mg/kg/day; n=9), or tangeretin (1.8 mg/kg/day; n=7) via osmotic minipumps for two weeks. The vigilance states were monitored in freely moving mice for 24 hours at baseline and on post-surgical day 14 upon spared nerve injury (SNI)^17^. Pain-like behaviour was assessed using von Frey filaments and brush dynamic tests at baseline and at post-surgical days 3, 7, and 14, after which the mice were euthanized and the ipsilateral DRGs were collected for rhythmicity monitoring in the bioluminescence device.

### Spared nerve injury surgery and minipump implantation

To induce NP, the animals underwent SNI surgery as described earlier^16^. Briefly, the tibial and common peroneal nerves of the sciatic nerve were ligated and dissected through 1-cm lateral left paw incision, while the sural nerve was left intact (“spared”). Sham surgery only exposed the sciatic nerve. Surgeries were performed under isoflurane anaesthesia, and animals received 10 mg/kg of carprofen (Rimadyl; Zoetis Animal Health, Finland) prior to the procedure. The muscle layer and skin incision were carefully closed with surgical sutures.

Directly after SNI surgeries, osmotic minipumps (Alzet model 1002; DURECT Corporation, USA) were implanted for continuous administration of flavonoids. Prior to implantation, pump loading was monitored by recording the weight of each osmotic minipump before and after addition of 10% DMSO in 90% sterile saline as vehicle, nobiletin, or tangeretin. Only pumps containing a solution volume exceeding 90% of the 101.5 µl capacity were used. Flow moderators were attached to the pumps as part of the preparation procedure. The pumps were carefully placed into the abdominal cavity via small (<1mm) skin and peritoneal wall incisions, which were subsequently closed with 6-0 surgical knots.

Povidone iodine was applied to the wound site for disinfection. Each animal received nobiletin (2 mg/kg/day), tangeretin (1.8 mg/kg/day) or vehicle, delivered continuously at a mean rate of 0.25 µl/h. The dosages were selected based on previously published studies^65,66^ and slightly modified according to pilot experiments assessing compound solubility in the vehicle for minipump delivery. Pump implantation was performed in a random order across the treatment groups to reduce batch effects. The animals’ activity, coat condition, and overall well-being were monitored for three days postoperatively to ensure adequate recovery from surgery.

### Behavioural testing

Behavioural testing, including von Frey filament and dynamic brush tests, was performed at baseline and on post-surgical days 3, 7, and 14 to assess mechanical hypersensitivity and dynamic allodynia. Testing was conducted in red acrylic cylinders placed on an elevated mesh platform. Each test began with a 30-min habituation period, with additional 1-h habituation periods conducted on two days before baseline testing. Behavioural tests were consistently performed within the same daytime window (ZT3– 6; lights on at 06:00) to minimise circadian variation.

### von Frey filament test

The filaments, starting at a force of 0.02 g, were applied at a 90-degree angle to the lateral region of the paw, corresponding to the region innervated by the spared sural nerve^67^. The force was gradually increased (0.04, 0.07, 0.16, 0.4, 0.6, 1.0) until three pain responses out of five filament applications were observed. The lowest force producing this response was defined as the paw withdrawal threshold. Positive pain responses included rapid paw withdrawal, shaking movements, or licking of the paw.

### Dynamic brush test

Dynamic allodynia was assessed by gently brushing the lateral surface of the hind paw with a paintbrush^68^. The responses were scored on a scale from 0 to 3, where 0 indicates no pain and 3 indicates a strong nociceptive response. Score 0 was characterized by a rapid lifting of the paw lasting less than 1 s. Pain responses included elevation of brushed paw towards the body for more than 2 s (score of 1), a lateral kick or a sudden jump-like response (score of 2) and repetitive flinching or licking of the brushed paw (score of 3).

### EEG/EMG transmitter implantation and recording

Transmitter implantation surgeries (HDX02, Data Sciences International, USA) were performed in randomized order under 1.5-2.5% isoflurane anaesthesia (Attane Vet; Piramal Critical Care B.V., The Netherlands), 14 days before SNI or sham surgeries. First, the mice were secured in a stereotaxic frame and the skin overlying the skull was opened to expose the surgical site. Two tiny openings were then drilled into the skull over the left frontal and parietal regions for placement of the EEG leads. The frontal biopotential lead was placed 1.0 mm anterior to bregma and 0.5 mm lateral to the midline, and the parietal lead 3.4 mm posterior to bregma and 1.5 mm lateral to the midline. Following placement of the EEG leads, dental cement (GC FujiCEM 2; Plandent, Finland) was applied to secure biopotential leads to the skull. EMG activity was measured from the cervical trapezius muscle using a pair of leads threaded through the muscle with a 20-gauge needle. The dental cement was allowed to set before closing the surgical wound with 6-0 coated Vicryl Plus sutures (Ethicon, USA). For perioperative analgesia, mice received carprofen (Rimadyl, Zoetis Animal Health) before surgery (10 mg/kg) and for two days postoperatively (5 mg/kg). The local anaesthetic lidocaine was applied to the wound site.

EEG/EMG signals were recorded by telemetry using PhysioTel receiver plates (RPC-1; Data Sciences International) placed under each cage. The telemetry setup was operated through a DSI Matrix 2.0 and signals were sampled at 500 Hz with a bandwidth of 0.5–80 Hz. Telemetry data were acquired in Spike2 (version 10.24; Cambridge Electronic Design, United Kingdom) through the DSI Talker interface. Two female mice from the sham group were excluded from the study, due to transmitter failure and surgical complications, resulting in the final group sizes of 5 SNI and 3 female sham mice.

### Sleep scoring and epoch count analysis

The sleep stages were manually scored in Spike2 using sleepscore_v1.86.s2s script (Cambridge Electronic Design) as in our previous study^16^. To avoid bias, all files were blinded and coded by an independent researcher who was not involved in sleep scoring or data analysis. Epochs were categorized as wake, NREM, REM or noise based on EEG frequency and amplitude together with EMG activity. High EMG activity combined with mixed-frequency EEG was assigned to wakefulness, while delta activity between 0.5 and 4 Hz with reduced EMG activity was scored as NREM sleep. REM sleep was characterized by dominant theta activity (5–9 Hz) accompanied by minimal EMG activity ^69^. The percentages of time spent in the different vigilance states are presented for all groups in supplementary Table 3. The number of epochs in each stage across the 24-h period was normalized to baseline to account for interindividual variability between animals. For each vigilance state, epoch counts were analysed over time by comparing baseline with post-surgical day 14 within each group, and between the SNI and sham groups on day 14.

### Sleep spindle analysis and sleep-wake transitions

Total number and duration of sleep spindles, along with sleep-wake transitions between the vigilance states were identified using an automated MATLAB algorithm (spindle_script_revised_17_5s) as previously described^16,32^. Briefly, signals were filtered within the sigma frequency range (10–15 Hz), and spindle events were identified using root mean square (RMS)-based detection calculated over 750-ms intervals. These values were raised to the third power to increase the contrast between spindle-related activity and background EEG. Spindle events were then detected using lower and upper thresholds of 1.2 and 3.5 times the average signal value across the whole recording. The algorithm also automatically quantified transitions between wakefulness, NREM sleep, and REM sleep.

### Power spectra analysis

Power spectral analysis was performed as previously described^16^. Briefly, the power spectra were calculated separately for each vigilance state using Spike2 software and fast Fourier transform with a 1,024-point size and a Hanning window. Data from the 12-h light and 12-h dark periods were evaluated independently. Statistical comparisons were performed at 0.5-Hz intervals across vigilance state-specific frequency ranges and at 5-Hz intervals across the 30–50 Hz low-gamma range, as shown in the Supplementary tables 1 and 2.

### Bioluminescence recordings

Upon completion of behavioural testing at post-surgical day 14, PER2::LUC mice were euthanized by cervical dislocation under isoflurane-anaesthesia and ipsilateral L3-L5 DRGs were dissected between ZT6 and ZT10. DRGs were immediately placed into ice-cold Hank’s buffer (Thermo Fisher Scientific, Waltham, MA, USA) and transported to the laboratory within 1 h of euthanasia. Harvested DRGs were transferred into a Petri dish containing 1.2ml of prewarmed Dulbecco’s Phosphate Buffered Saline (pH 7.2) supplemented with 10 mM HEPES, 2% B-27, 5 U/ml penicillin and 5 μg/ml streptomycin as well as 0.2 mM D-luciferin potassium salt (Thermo Fisher), under dim red light (Model IL21, Beurer GmbH, Ulm, Germany) to minimize light-induced circadian disruption.

The sealed dishes were placed in a LumiCycle 32 bioluminescence device (Actimetrics, Lafayette, IN, USA), kept in a 36°C non-humidified incubator, and photon counts representing *Per2* rhythmicity were recorded for each sample every 10 min for 6 days. The *Per2* signals (counts/s) were normalized in a LumiCycle Analysis software (version 3.103; Actimetrics, USA) by applying a 24-h running average baseline subtraction, with a 1-hour moving average filtering. Three circadian parameters, amplitude, (the distance between the peak and midline), period (the duration of one oscillation) and acrophase (the point when the rhythm reaches its first peak), were analysed from damped sine-wave fits using LM Fit^70^. Rhythmicity was assessed across three complete oscillations and further evaluated by relative rhythmic power using periodogram analysis. Bioluminescence data was filtered for quality control based on baseline stability and curve-fitting convergence (goodness-of-fit typically >75%) before rhythmicity analysis.

### Statistics

Changes in behavioural outcomes were assessed by two-way analysis of variance (ANOVA), followed by Tukey’s tests for multiple comparisons. Group differences were analysed using two-tailed Student’s unpaired t-tests for baseline-normalized epoch counts and spindle measures at post-surgical day 14, as well as for sex differences in baseline epoch counts. Prior to the t-test, data normality was assessed using the Shapiro–Wilk test. If the data were not normally distributed, the non-parametric Mann– Whitney test was used instead. Baseline and day 14 values within each group were compared using two-tailed Student’s paired t-test. For power spectra, group differences at baseline and post-surgical day 14 were analysed using two-tailed Student’s unpaired t-tests. One-way ANOVA followed by Tukey’s post hoc test was carried out to evaluate differences in circadian parameters. All analyses were performed and figures were created using GraphPad Prism version 10.1.2 (Boston, MA, USA) and BioRender (Toronto, Canada). A p-value ≤ 0.05 was deemed statistically significant and behavioural results are reported as mean ± SEM.

## Contributions

Conceptualisation: V.P., Analysis: H.L., W.D., M.W., Visualisation: H.L., Writing original draft: H.L., Writing review & editing: H.L., W.D., M.W., N.M., MK.K., A.P., E.K., V.P., Methodology: H.L., W.D., M.W., N.M., MK.K., HK.W, Resources: A.P., HK.W., Funding acquisition: V.P., E.K., Supervision: V.P.

