## Supplemental File for "Flavonoid chronotherapy normalizes circadian dynamics and sleep architecture in mice with painful nerve injury"

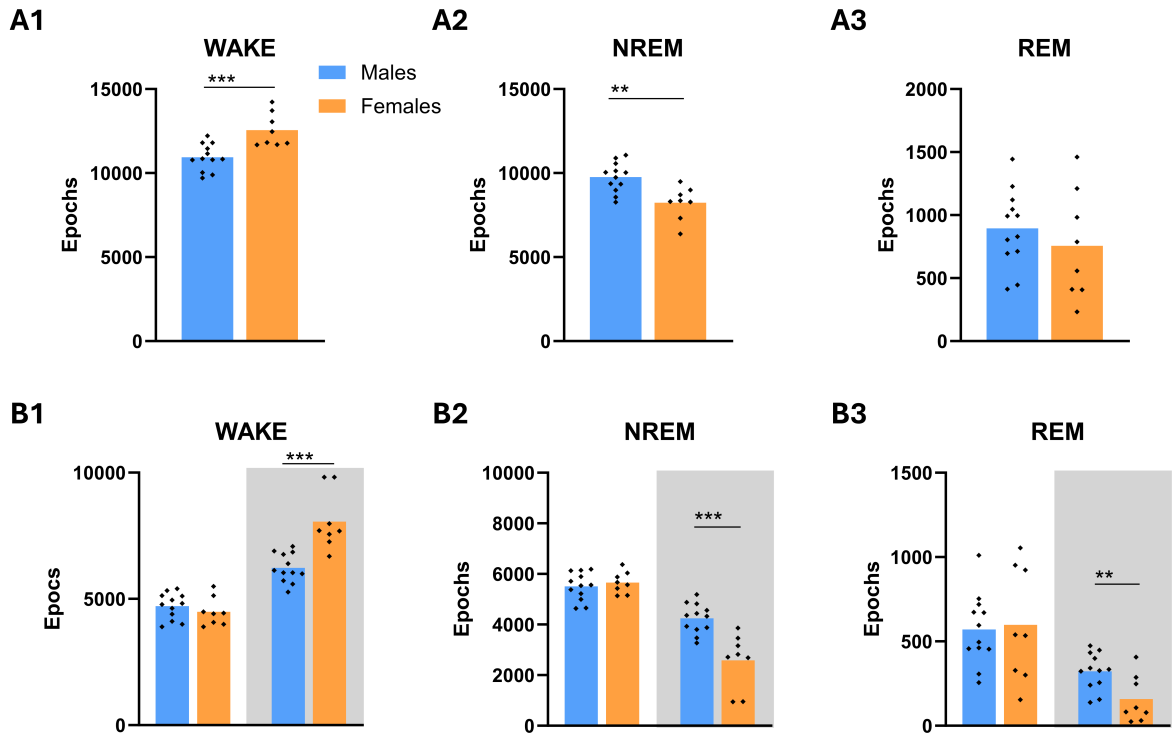

Supplementary Figure 1. **Baseline sex differences in sleep architecture of PER2::LUC mice.** **A)** Comparison of 24-h sleep-wake epochs (4 s) between male (n=12) and female (n=8) PER2::LUC mice during wake, non-rapid eye movement (NREM) sleep, and rapid eye movement (REM) sleep. Females showed fewer NREM (A2) epochs and more wake epochs than males (A1), while REM epochs remained unchanged (A3). **B)** Sleep-wake states assessed separately across the light and dark periods. During the dark period, females exhibited increased wakefulness (B1) and reduced NREM (B2) and REM sleep (B3) compared with males. \*\*p<0.01, \*\*\*p<0.001.

Supplementary Table 1. **Significant frequencies identified in EEG power spectra analyses during the light phase, corresponding to Figure 6.** Spectral power was assessed at 0.5-Hz resolution from 0.5 to 6 Hz in wake (**A**), from 0.5 to 10 Hz in NREM (**B**), and from 0.5 to 9.5 Hz in REM (**C**), and within the 30–50 Hz low-gamma band at 5-Hz resolution, both at baseline and on day 14. The table includes frequencies showing significant effects on post-surgical day 14 that were absent at baseline, as well as significant within-group differences between baseline and day 14. ns= non-significant, SNI= spared nerve injury, Hz= hertz, BL= baseline.

| A |  | WAKE |  |
| --- | --- | --- | --- |
| BL vs. day 14 | Sham | 0.5-6 Hz | 30-50 Hz |
|  | SNI vehicle | ns. | ns. |
|  | SNI + nobiletin | ns. | 30 Hz: t(6) = 3.2, p = 0.018<br>35 Hz: t(6) = 2.7, p = 0.034<br>50 Hz: t(6) = 2.5, p = 0.045 |
|  | SNI+ tangeretin | ns. | 40 Hz: t(6) = 2.7, p = 0.038 |
| Day 14 | Sham vs. SNI vehicle | 6 Hz: t(10) = 2.5, p = 0.032 |  |
|  | Sham vs. SNI+ nobiletin | ns. |  |
|  | Sham vs. SNI+ tangeretin | ns. |  |
|  | SNI vehicle<br>vs.<br>SNI +nobiletin | ns. | 30 Hz: t(14) = 2.2, p = 0.046<br>50 Hz: t(14) = 2.5, p = 0.026 |
|  | SNI vehicle<br>vs.<br>SNI+ tangeretin | ns. |  |
|  | SNI+ nobiletin<br>vs.<br>SNI+ tangeretin | 1 Hz: t(10) = 2.6, p = 0.023<br>1.5 Hz: t(10) = 2.5, p = 0.024<br>2 Hz: t(10) = 2.4, p = 0.031<br>2.5 Hz: t(10) = 2.2, p = 0.043 |  |

## B

### NREM

|  |  | 0.5-10 Hz | 30-50 Hz |
| --- | --- | --- | --- |
| **BL**<br>**vs.**<br>**day 14** | Sham | ns. | ns. |
|  | SNI vehicle | 3 Hz: t(6) = 2.7, p = 0.035<br>3.5 Hz: t(6) = 3.6, p = 0.012<br>4.5 Hz: t(6) = 3.2, p = 0.018<br>5 Hz: t(6) = 4.9, p = 0.003<br>5.5 Hz: t(6) = 5.5, p = 0.002<br>6 Hz: t(6) = 5.1, p = 0.002<br>6.5 Hz: t(6) = 4.7, p = 0.003<br>7 Hz: t(6) = 4.4, p = 0.004<br>7.5 Hz: t(6) = 4.3, p = 0.005<br>8 Hz: t(6) = 4.3, p = 0.005<br>8.5 Hz: t(6) = 3.9, p = 0.008<br>9 Hz: t(6) = 4.1, p = 0.007<br>9.5 Hz: t(6) = 3.9, p = 0.008<br>10 Hz: t(6) = 3.8, p = 0.009 | 30 Hz: t(6) = 4.0, p = 0.008<br>35 Hz: t(6) = 3.9, p = 0.008<br>40 Hz: t(6) = 4.1, p = 0.006<br>45 Hz: t(6) = 3.7, p = 0.010<br>50 Hz: t(6) = 3.6, p = 0.012 |
|  | SNI+ nobiletin | 2 Hz: t(8) = 2.8, p = 0.023<br>2.5 Hz: t(8) = 3.4, p = 0.009<br>4.5 Hz: t(8) = 3.2, p = 0.013<br>5 Hz: t(8) = 3.7, p = 0.006<br>5.5 Hz: t(8) = 3.7, p = 0.006<br>6 Hz: t(8) = 3.3, p = 0.010<br>6.5 Hz: t(8) = 3.0, p = 0.016<br>7 Hz: t(8) = 2.9, p = 0.021<br>7.5 Hz: t(8) = 2.7, p = 0.025<br>8 Hz: t(8) = 2.7, p = 0.027<br>8.5 Hz: t(8) = 2.7, p = 0.026<br>9 Hz: t(8) = 2.8, p = 0.023<br>9.5 Hz: t(8) = 3.0, p = 0.018<br>10 Hz: t(8) = 3.2, p = 0.013 | 30 Hz: t(8) = 4.5, p = 0.002<br>35 Hz: t(8) = 4.2, p = 0.003<br>40 Hz: t(8) = 4.0, p = 0.004<br>45 Hz: t(8) = 4.3, p = 0.003<br>50 Hz: t(8) = 3.8, p = 0.006 |
|  | SNI+ tangeretin | 2.5 Hz: t(6) = 3.1, p = 0.020<br>3 Hz: t(6) = 3.5, p = 0.013<br>4 Hz: t(6) = 2.7, p = 0.037<br>4.5 Hz: t(6) = 4.1, p = 0.006<br>5 Hz: t(6) = 3.6, p = 0.011<br>5.5 Hz: t(6) = 3.3, p = 0.017<br>6 Hz: t(6) = 3.4, p = 0.015<br>6.5 Hz: t(6) = 3.5, p = 0.013<br>7 Hz: t(6) = 3.6, p = 0.012<br>7.5 Hz: t(6) = 3.3, p = 0.017<br>8 Hz: t(6) = 2.5, p = 0.044<br>8.5 Hz: t(6) = 2.5, p = 0.044<br>9 Hz: t(6) = 2.7, p = 0.036<br>9.5 Hz: t(6) = 2.6, p = 0.039<br>10 Hz: t(6) = 2.5, p = 0.045 | 30 Hz: t(6) = 4.3, p = 0.005<br>35 Hz: t(6) = 5.3, p = 0.002<br>40 Hz: t(6) = 5.8, p = 0.001<br>45 Hz: t(6) = 5.0, p = 0.002<br>50 Hz: t(6) = 3.8, p = 0.010 |
| **Day 14** | Sham vs. SNI | 3 Hz: t(10) = 2.4, p = 0.034<br>3.5 Hz: t(10) = 2.7, p = 0.022<br>5 Hz: t(10) = 2.5, p = 0.030<br>5.5 Hz: t(10) = 2.8, p = 0.020 | ns. |
| | | 6 Hz: $t(10) = 3.0$, $p = 0.014$<br>6.5 Hz: $t(10) = 3.1$, $p = 0.011$ | |
|  | Sham<br>vs.<br>SNI+ nobiletin | ns. | ns. |
|  | Sham<br>vs.<br>SNI+ tangeretin | ns. | ns. |
| | SNI vehicle<br>vs.<br>SNI +nobiletin | 0.5 Hz: $t(14) = 2.5$, $p = 0.027$<br>1 Hz: $t(14) = 2.3$, $p = 0.035$<br>1.5 Hz: $t(14) = 2.4$, $p = 0.030$<br>2 Hz: $t(14) = 2.7$, $p = 0.017$<br>3.5 Hz: $t(14) = 3.8$, $p = 0.002$<br>4 Hz: $t(14) = 3.1$, $p = 0.008$<br>9.5 Hz: $t(14) = 2.2$, $p = 0.045$<br>10 Hz: $t(14) = 2.2$, $p = 0.046$ | 30 Hz: $t(14) = 2.4$, $p = 0.032$<br>35 Hz: $t(14) = 2.5$, $p = 0.025$<br>40 Hz: $t(14) = 2.4$, $p = 0.030$<br>45 Hz: $t(14) = 2.3$, $p = 0.035$<br>50 Hz: $t(14) = 2.2$, $p = 0.044$ |
| | SNI vehicle<br>vs.<br>SNI+ tangeretin | 8.5 Hz: $t(12) = 2.3$, $p = 0.040$<br>9 Hz: $t(12) = 2.4$, $p = 0.034$<br>9.5 Hz: $t(12) = 2.4$, $p = 0.035$<br>10 Hz: $t(12) = 2.3$, $p = 0.041$ | ns. |
|  | SNI+ nobiletin<br>vs.<br>SNI+ tangeretin |  | ns. |

**C**

**REM**

|  |  |  |  |
| --- | --- | --- | --- |
|  |  | **3-9.5 Hz** | **30-50 Hz** |
| **BL**<br>**vs.**<br>**day 14** | Sham | ns. | ns. |
| | SNI vehicle | 3 Hz: $t(6) = 3.9$, $p = 0.009$<br>3.5 Hz: $t(6) = 2.6$, $p = 0.042$<br>4 Hz: $t(6) = 2.6$, $p = 0.043$<br>4.5 Hz: $t(6) = 4.6$, $p = 0.004$<br>5 Hz: $t(6) = 5.4$, $p = 0.002$<br>5.5 Hz: $t(6) = 4.7$, $p = 0.003$<br>6 Hz: $t(6) = 3.8$, $p = 0.009$<br>6.5 Hz: $t(6) = 5.0$, $p = 0.002$<br>7 Hz: $t(6) = 5.1$, $p = 0.002$<br>7.5 Hz: $t(6) = 4.3$, $p = 0.005$<br>8 Hz: $t(6) = 4.9$, $p = 0.003$<br>8.5 Hz: $t(6) = 4.4$, $p = 0.005$<br>9 Hz: $t(6) = 5.3$, $p = 0.002$<br>9.5 Hz: $t(6) = 6.1$, $p = 0.0009$ | 30 Hz: $t(6) = 4.6$, $p = 0.004$<br>35 Hz: $t(6) = 4.4$, $p = 0.005$<br>40 Hz: $t(6) = 3.5$, $p = 0.013$<br>45 Hz: $t(6) = 4.7$, $p = 0.003$<br>50 Hz: $t(6) = 3.2$, $p = 0.020$ |
| | SNI+ nobiletin | 6 Hz: $t(8) = 2.4$, $p = 0.046$<br>6.5 Hz: $t(8) = 2.4$, $p = 0.045$<br>7.5 Hz: $t(8) = 2.4$, $p = 0.046$<br>8 Hz: $t(8) = 2.7$, $p = 0.028$<br>8.5 Hz: $t(8) = 2.3$, $p = 0.049$ | 30 Hz: $t(8) = 5.1$, $p = 0.00098$<br>35 Hz: $t(8) = 5.1$, $p = 0.00098$<br>40 Hz: $t(8) = 5.2$, $p = 0.00081$<br>45 Hz: $t(8) = 4.5$, $p = 0.002$<br>50 Hz: $t(8) = 3.8$, $p = 0.005$ |
| | SNI+ tangeretin | ns. | 30 Hz: $t(6) = 5.0$, $p = 0.002$<br>35 Hz: $t(6) = 3.7$, $p = 0.00985$<br>40 Hz: $t(6) = 5.0$, $p = 0.002$<br>45 Hz: $t(6) = 3.8$, $p = 0.009$ |
| **Day 14** | Sham vs. SNI vehicle | 3 Hz: $t(10) = 2.4$, $p = 0.038$<br>5.5 Hz: $t(10) = 3.1$, $p = 0.0104$<br>6 Hz: $t(10) = 3.0$, $p = 0.013$<br>6.5 Hz: $t(10) = 3.4$, $p = 0.007$<br>7 Hz: $t(10) = 3.4$, $p = 0.0068$ | ns. |
|  | Sham<br>vs.<br>SNI+ nobiletin | ns. | ns. |
|  | Sham<br>vs.<br>SNI+ tangeretin | ns. | ns. |
| | SNI vehicle<br>vs.<br>SNI +nobiletin | 3 Hz: $t(14) = 2.3, p = 0.041$ | ns. |
| | SNI vehicle<br>vs.<br>SNI+ tangeretin | 7.5 Hz: $t(12) = 2.3, p = 0.039$<br>8 Hz: $t(12) = 3.0, p = 0.012$<br>8.5 Hz: $t(12) = 2.9, p = 0.013$<br>9 Hz: $t(12) = 3.0, p = 0.011$<br>9.5 Hz: $t(12) = 3.0, p = 0.011$ | ns. |
|  | SNI+ nobiletin<br>vs.<br>SNI+ tangeretin | ns. | ns. |

Supplementary Table 2. **Significant EEG power spectrum frequencies observed during the dark phase and shown in Figure 7.** EEG spectral power was evaluated at 0.5-Hz resolution across 0.5–6 Hz in wake (A), 0.5–10 Hz in NREM (B), and 0.5–9.5 Hz in REM (C). The 30–50 Hz low-gamma range was analyzed separately at 5-Hz resolution. The analyses were conducted at baseline and on day 14. The table reports frequencies that showed significant effects on post-surgical day 14 but not at baseline, as well as significant changes within groups between baseline and day 14. ns = non-significant, SNI = spared nerve injury, Hz = hertz, BL = baseline.

**A**

**WAKE**

|  |  | 0.5-6 Hz | 30-50 Hz |
| --- | --- | --- | --- |
| **BL**<br>**vs.**<br>**day 14** | Sham | ns. | ns. |
| | SNI vehicle | 6 Hz: $t(6) = 2.7, p = 0.038$ | 30 Hz: $t(6) = 4.9, p = 0.003$<br>35 Hz: $t(6) = 4.1, p = 0.006$<br>40 Hz: $t(6) = 3.8, p = 0.009$<br>45 Hz: $t(6) = 3.5, p = 0.013$<br>50 Hz: $t(6) = 3.7, p = 0.010$ |
| | SNI+ nobiletin | ns. | 30 Hz: $t(6) = 4.9, p = 0.003$<br>35 Hz: $t(6) = 4.1, p = 0.006$<br>40 Hz: $t(6) = 3.8, p = 0.009$<br>45 Hz: $t(6) = 3.5, p = 0.013$<br>50 Hz: $t(6) = 3.7, p = 0.010$ |
| | SNI+ tangeretin | ns. | 35 Hz: $t(6) = 2.5, p = 0.046$ |
| **Day 14** | Sham<br>vs.<br>SNI vehicle | 5 Hz: $t(10) = 2.3, p = 0.047$<br>5.5 Hz: $t(10) = 2.5, p = 0.032$<br>6 Hz: $t(10) = 2.9, p = 0.015$ | ns. |
| | Sham<br>vs.<br>SNI+ nobiletin | 4 Hz: $t(12) = 2.3, p = 0.040$<br>6 Hz: $t(12) = 2.4, p = 0.031$ | ns. |
|  | Sham<br>vs.<br>SNI+ tangeretin |  | ns. |
| | SNI vehicle<br>vs.<br>SNI +nobiletin | | 50 Hz: $t(14) = 2.4, p = 0.030$ |
|  | SNI vehicle<br>vs.<br>SNI+ tangeretin |  | ns. |
| | SNI+ nobiletin<br>vs.<br>SNI+ tangeretin | 1 Hz: $t(14) = 2.2, p = 0.043$ | ns. |

**B**

**NREM**

|  |  | **0.5-10 Hz** | **30-50 Hz** |
| --- | --- | --- | --- |
| **BL**<br>**vs.**<br>**day 14** | Sham | ns. | ns. |
|  | SNI vehicle | 1 Hz: t(6) = 2.5, p = 0.049<br>1.5 Hz: t(6) = 3.2, p = 0.018<br>2 Hz: t(6) = 3.3, p = 0.016<br>2.5 Hz: t(6) = 2.7, p = 0.036<br>3 Hz: t(6) = 2.6, p = 0.041<br>3.5 Hz: t(6) = 5.1, p = 0.002<br>4 Hz: t(6) = 4.3, p = 0.005<br>4.5 Hz: t(6) = 3.5, p = 0.014<br>5 Hz: t(6) = 6.2, p = 0.00082<br>5.5 Hz: t(6) = 8.2, p = 0.00018<br>6 Hz: t(6) = 8.0, p = 0.00020<br>6.5 Hz: t(6) = 7.9, p = 0.00022<br>7 Hz: t(6) = 7.6, p = 0.00026<br>7.5 Hz: t(6) = 7.3, p = 0.00034<br>8 Hz: t(6) = 5.8, p = 0.00118<br>8.5 Hz: t(6) = 5.3, p = 0.002<br>9 Hz: t(6) = 4.7, p = 0.003<br>9.5 Hz: t(6) = 4.4, p = 0.004<br>10 Hz: t(6) = 4.9, p = 0.003 | 30 Hz: t(6) = 4.5, p = 0.004<br>35 Hz: t(6) = 5.0, p = 0.003<br>40 Hz: t(6) = 4.6, p = 0.004<br>45 Hz: t(6) = 4.2, p = 0.005<br>50 Hz: t(6) = 3.7, p = 0.010 |
|  | SNI+ nobiletin | 2 Hz: t(8) = 2.5, p = 0.036<br>2.5 Hz: t(8) = 4.1, p = 0.003<br>3 Hz: t(8) = 4.7, p = 0.002<br>3.5 Hz: t(8) = 3.6, p = 0.008<br>5 Hz: t(8) = 3.7, p = 0.007<br>5.5 Hz: t(8) = 4.4, p = 0.002<br>6 Hz: t(8) = 4.5, p = 0.002<br>6.5 Hz: t(8) = 4.5, p = 0.002<br>7 Hz: t(8) = 4.4, p = 0.002<br>7.5 Hz: t(8) = 3.8, p = 0.005<br>8 Hz: t(8) = 3.1, p = 0.014<br>8.5 Hz: t(8) = 2.9, p = 0.018<br>9 Hz: t(8) = 3.1, p = 0.015<br>9.5 Hz: t(8) = 3.1, p = 0.016<br>10 Hz: t(8) = 3.2, p = 0.013 | 30 Hz: t(8) = 4.2, p = 0.003<br>35 Hz: t(8) = 4.5, p = 0.002<br>40 Hz: t(8) = 4.6, p = 0.002<br>45 Hz: t(8) = 4.3, p = 0.003<br>50 Hz: t(8) = 4.0, p = 0.004 |
|  | SNI+ tangeretin | 1.5 Hz: t(6) = 2.6, p = 0.041<br>2 Hz: t(6) = 3.4, p = 0.015<br>2.5 Hz: t(6) = 4.1, p = 0.006<br>3 Hz: t(6) = 2.9, p = 0.027<br>5.5 Hz: t(6) = 3.3, p = 0.016<br>6 Hz: t(6) = 3.5, p = 0.013<br>6.5 Hz: t(6) = 3.4, p = 0.015<br>7 Hz: t(6) = 2.5, p = 0.047 | 35 Hz: t(6) = 2.7, p = 0.036 |
|  | Sham<br>vs.<br>SNI vehicle | 3 Hz: t(10) = 2.4, p = 0.039<br>5 Hz: t(10) = 2.7, p = 0.022<br>5.5 Hz: t(10) = 2.9, p = 0.015<br>6 Hz: t(10) = 3.2, p = 0.010 | ns. |
|  | Sham<br>vs.<br>SNI+ nobiletin | ns. | ns. |
| **Day 14** | Sham<br>vs.<br>SNI+ tangeretin | ns. | 50 Hz: t(10) = 2.3, p = 0.046 |
|  | SNI vehicle<br>vs.<br>SNI +nobiletin | 0.5 Hz: t(14) = 2.3, p = 0.034<br>1 Hz: t(14) = 2.3, p = 0.035<br>1.5 Hz: t(14) = 2.3, p = 0.036<br>3.5 Hz: t(14) = 3.2, p = 0.006<br>4 Hz: t(14) = 3.0, p = 0.009 | 35 Hz: t(14) = 2.5, p = 0.023<br>40 Hz: t(14) = 2.5, p = 0.025<br>45 Hz: t(14) = 2.4, p = 0.030<br>50 Hz: t(14) = 2.3, p = 0.040 |
|  | SNI vehicle<br>vs. | 7.5 Hz: t(12) = 2.4, p = 0.033<br>8 Hz: t(12) = 2.5, p = 0.031 | ns. |
|  | SNI+ tangeretin | 8.5 Hz: t(12) = 2.5, p = 0.028<br>9 Hz: t(12) = 2.5, p = 0.029<br>9.5 Hz: t(12) = 2.5, p = 0.027<br>10 Hz: t(12) = 2.5, p = 0.026 |  |
|  | SNI+ nobiletin<br>vs.<br>SNI+ tangeretin | ns. | ns. |

**C**

**REM**

|  |  | **3-9.5 Hz** | **30-50 Hz** |
| --- | --- | --- | --- |
| **BL**<br>**vs.**<br>**day 14** | Sham | ns. | ns. |
|  | SNI vehicle | 3 Hz: t(6) = 5.0, p = 0.002<br>3.5 Hz: t(6) = 4.0, p = 0.007<br>7 Hz: t(6) = 3.5, p = 0.012<br>7.5 Hz: t(6) = 10.8, p = 0.00004<br>8 Hz: t(6) = 10.4, p = 0.00005<br>8.5 Hz: t(6) = 8.1, p = 0.0002<br>9 Hz: t(6) = 7.4, p = 0.0003<br>9.5 Hz: t(6) = 8.1, p = 0.0002 | 30 Hz: t(6) = 7.1, p = 0.0004<br>35 Hz: t(6) = 7.6, p = 0.0003<br>40 Hz: t(6) = 7.0, p = 0.0004<br>45 Hz: t(6) = 6.1, p = 0.0009<br>50 Hz: t(6) = 6.6, p = 0.0006 |
|  | SNI+ nobiletin | 3 Hz: t(8) = 3.1, p = 0.016<br>3.5 Hz: t(8) = 3.3, p = 0.011<br>5 Hz: t(8) = 2.7, p = 0.025<br>5.5 Hz: t(8) = 2.4, p = 0.046<br>7.5 Hz: t(8) = 2.8, p = 0.023<br>8 Hz: t(8) = 2.9, p = 0.021<br>8.5 Hz: t(8) = 2.5, p = 0.037<br>9.5 Hz: t(8) = 2.5, p = 0.037 | 30 Hz: t(8) = 5.6, p = 0.0005<br>35 Hz: t(8) = 5.4, p = 0.0006<br>40 Hz: t(8) = 5.8, p = 0.0004<br>45 Hz: t(8) = 5.4, p = 0.0007<br>50 Hz: t(8) = 4.7, p = 0.002 |
|  | SNI+ tangeretin | 6 Hz: t(5) = 3.1, p = 0.026<br>6.5 Hz: t(5) = 4.5, p = 0.007<br>7 Hz: t(5) = 3.9, p = 0.012 | 35 Hz: t(6) = 3.5, p = 0.013<br>50 Hz: t(6) = 2.6, p = 0.039 |
| **Day 14** | Sham<br>vs.<br>SNI vehicle | 3 Hz: t(10) = 2.6, p = 0.029<br>3.5 Hz: t(10) = 2.3, p = 0.046<br>8 Hz: t(10) = 4.0, p = 0.002<br>8.5 Hz: t(10) = 3.9, p = 0.003<br>9 Hz: t(10) = 3.9, p = 0.003 | ns. |
|  | Sham<br>vs.<br>SNI+ nobiletin | ns. | ns. |
|  | Sham<br>vs.<br>SNI+ tangeretin | ns. | ns. |
|  | SNI vehicle<br>vs.<br>SNI +nob | 3 Hz: t(14) = 2.2, p = 0.047 | ns. |
|  | SNI vehicle<br>vs.<br>SNI+ tangeretin | 7.5 Hz: t(12) = 2.3, p = 0.043<br>8 Hz: t(12) = 3.0, p = 0.011<br>8.5 Hz: t(12) = 3.2, p = 0.007<br>9 Hz: t(12) = 3.5, p = 0.004<br>9.5 Hz: t(12) = 3.6, p = 0.003 | 30 Hz: t(12) = 2.3, p = 0.037<br>35 Hz: t(12) = 2.6, p = 0.022<br>40 Hz: t(12) = 2.4, p = 0.031<br>45 Hz: t(12) = 2.2, p = 0.04987 |
|  | SNI+ nobiletin<br>vs.<br>SNI+ tangeretin | ns. | ns. |

Supplementary Table 3. **Percentage of recording time spent in each vigilance state in each group at baseline (A) and postsurgical day 14 (B).** Values represent the percentage of total scored epochs classified as wake, non-rapid eye movement (NREM), rapid eye movement (REM), or noise for each experimental group and are presented as mean ± SEM.

| **A** | **Sham**<br>**male** | **Sham**<br>**female** | **SNI**<br>**male** | **SNI**<br>**female** | **SNI+nobiletin**<br>**male** | **SNI+ tangeretin**<br>**male** |
| --- | --- | --- | --- | --- | --- | --- |
| **Wake (%)** | 49.2 ± 1.7 | 54.3 ± 0.2 | 51.7 ± 1.3 | 60.4 ± 2.0 | 51.2 ± 1.2 | 49.2 ± 1.7 |
| **NREM (%)** | 47.1 ± 1.4 | 42.0 ± 1.1 | 43.7 ± 1.6 | 35.8 ± 1.8 | 44.2 ± 1.2 | 47.0 ± 1.4 |
| **REM (%)** | 3.7 ± 0.5 | 3.7 ± 1.1 | 4.5 ± 0.6 | 3.4 ± 1.0 | 4.5 ± 0.5 | 3.8 ± 0.7 |
| **Noise (%)** | 0 ± 0 | 0 ± 0 | 0.1 ± 0.08 | 0.4 ± 0.4 | 0.01 ± 0.01 | 0 ± 0 |

| **B** | **Sham**<br>**male** | **Sham**<br>**female** | **SNI**<br>**male** | **SNI**<br>**female** | **SNI+Nobiletin**<br>**male** | **SNI+ Tangeretin**<br>**male** |
| --- | --- | --- | --- | --- | --- | --- |
| **Wake (%)** | 48.0 ± 1.7 | 54.3 ± 1.5 | 45.0 ± 1.0 | 51.8 ± 3.8 | 46.4 ± 0.8 | 47.4 ± 1.0 |
| **NREM (%)** | 49.1 ± 1.4 | 42.6 ± 2.4 | 50.2 ± 1.1 | 39.9 ± 1.9 | 48.6 ± 1.0 | 48.8 ± 0.5 |
| **REM (%)** | 2.9 ± 0.7 | 3.2 ± 1.4 | 4.6 ± 0.6 | 3.3 ± 1.1 | 5.0 ± 0.7 | 3.8 ± 0.6 |
| **Noise (%)** | 0 ± 0 | 0 ± 0 | 0.3 ± 0.2 | 5.0 ± 4.8 | 0.1 ± 0.1 | 0.03 ± 0.03 |
